# Locus Coeruleus to Medial Prefrontal Cortex Noradrenergic Neural Circuit Modulates States of Consciousness during Sevoflurane Anesthesia in Mice

**DOI:** 10.1101/2025.05.01.651682

**Authors:** WeiHui Shao, Yue Yang, YaXuan Wu, XuanYi Di, ZiWen Zhang, Lu Liu, LeYuan Gu, XiaoXia Xu, ZhuoYue Zhang, JiaXuan Gu, HongHai Zhang

**Author notes:** Corresponding authors: HongHai Zhang. These authors contributed equally to this work (WeiHui Shao, Yue Yang, YaXuan Wu, XuanYi Di, ZiWen Zhang).

## Abstract

How exactly does general anesthesia achieve its effects? It is so widely used in surgeries and medical treatments to enhance comfort for nearly one hundred years, yet its precise mechanisms underlying the loss and recovery of consciousness still remain unclear. Utilizing a variety of research methods-pharmacological, optogenetic, chemogenetic, fiber photometry, and gene knockdown approaches-our study has shed light on the significant modulatory function of locus coeruleus noradrenergic neurons during the transition from sevoflurane anesthesia to awakening. Furthermore, the activation of the LC-mPFC circuit has been found to have a substantial arousal effect, with α1-AR playing a crucial role in this process. Additionally, GABAA-R has been identified as the key binding site for sevoflurane within the locus coeruleus. These findings collectively offer novel insights into the neural network mechanisms underlying general anesthesia, advancing our understanding of this complex medical phenomenon.

## 1 Introduction

General anesthesia is employed to induce a temporary state during surgeries or comfort-focused treatments to ensure patient safety and comfort. It puts patients into a temporary dreamland where they don’t feel pain, their reflexes are tamed, and their muscles chill out^1^. This is all thanks to the anesthetic drugs that give the central nervous system a gentle but firm "time-out". Previously, the journal Science listed “how general anesthetic drugs work” as one of the 125 important and pressing frontier scientific questions, due to their widespread use despite the unclear mechanisms.^2^

The alteration of consciousness induced by general anesthetic drugs is the core of research into the mechanisms of general anesthesia.^3^ Given the numerous neurophysiological parallels between general anesthesia and natural sleep, researchers have increasingly turned their attention in recent years to understanding how general anesthetics specifically target sleep-wake nuclei and neural circuits, ultimately leading to altered states of consciousness.^4^ This line of inquiry has emerged as a vital key to unlocking the mysteries of how general anesthesia works.^5^

Among the diverse array of anesthetic agents, sevoflurane stands out because of its rapid onset of action, fast recovery, little irritation to the airways, and pretty stable vital signs keeping.^6^ No wonder it’s a favorite in operating rooms and clinics today. However, despite its widespread use, we still don’t know exactly how it makes people lose consciousness and then wake them up again. Figuring this question out is crucial, as it not only enhances the safety and precision of general anesthesia but also propels advancements in anesthesiology and the development of new anesthetic agents.

The locus coeruleus (LC) serves as the central hub in the brain for producing norepinephrine (NE).^7^ NE-producing neurons in the LC are vital for modulating sleep-wake transitions, attention, stress-related behaviors, stress responses, and pain signaling.^8^ The activity of LC^NE^ neurons is tightly connected with arousal levels. These neurons are highly active when we are awake, driving increased NE release. In contrast, during sleep, their activity drops, resulting in diminished NE release.^9^ What’s more, stimulating the LC can jolt the brain from sleep to wakefulness.^10, 11^ Recent studies have revealed that other key players in the brain—like the serotonergic system in the dorsal raphe nucleus (DRN)^12^, the glutamatergic system in the parabrachial nucleus (PBN)^13^, the histaminergic system in the tuberomammillary nucleus (TMN)^14^, and the dopaminergic system in the ventral tegmental area (VTA)^15^—all have a say in whether we’re asleep or awake. Thus, our initial step was to make sure whether LC^NE^ neurons are involved in regulating the anesthesia-wakefulness process under sevoflurane anesthesia.

The medial prefrontal cortex (mPFC) is a crucial component of the prefrontal cortex. The mPFC is a key player in the brain’s sleep-wake cycle regulation and is like the brain’s control center for alertness^16, 17^, which handles a bunch of important tasks, from thinking and feeling to keeping us focused and aware.^18–21^ When awake, the mPFC is super active, constantly processing what’s going on around us and putting it all together. But during sleep, especially in the non-rapid eye movement (NREM) phase, its activity drops significantly.^22^ While other brain regions, such as the visual cortex and areas involved in emotional regulation, show increased activity during rapid eye movement (REM) sleep, the activity of the mPFC still remains relatively low.^23^ Research using fMRI shows that if we don’t get enough sleep, the mPFC’s activity goes down even more.^24^ In addition, research has suggested that the mPFC may be the spot in the brain through which general anesthetics induce unconsciousness by influencing the integration of information.^25–27^ LC^NE^ neurons send out connections to many brain regions, creating a huge network that controls how alert we are and how conscious we feel. Notably, the mPFC, being one of the main projection areas of the LC, is probably super important in this whole brain network. Thus, we hypothesize that the LC-mPFC NE neurocircuitry may be the key in regulating the shifts in consciousness associated with general anesthesia and wakefulness.

The research has uncovered that sevoflurane initially functions by targeting the GABAA receptors (GABAA-R) within the locus coeruleus. We found that the LC-mPFC NEergic pathway is really important for waking up from sevoflurane anesthesia, and it mostly happens through the alpha-1 adrenergic receptor (α1-AR). Initially, we employed pharmacological methods to investigate the connection between the NE system and the emergence from sevoflurane administration. After that, we turned to more precise and targeted techniques, like optogenetics and chemogenetics. These methods helped us dig deeper and really nail down our findings. We were able to show, without a doubt, that the LC-mPFC NEergic pathway plays a key role in waking up from sevoflurane anesthesia. In addition, we tested both male and female mice in our chemogenetic activation experiments. What we found is that the same strategies that help male mice wake up from sevoflurane-induced unconsciousness work just as well for female mice.

Building on the findings from our earlier experiments, we used knockdown techniques to selectively tweak the GABAA-R on the LC^NE^ neurons. We found that knocking down the GABAA-R not only shortened the recovery time from sevoflurane anesthesia but also impacted the calcium signaling at the noradrenergic terminals in the mPFC. More importantly, specifically blocking the α1-AR in the mPFC can reverse the shortened recovery time caused by knocking down the GABAA-R on LC^NE^ neurons.

For this research, we pulled out all the stops, using a mix of pharmacology, optogenetics, chemogenetics, calcium fiber photometry, and viral tracing to put our hypothesis to the test. Our study has shown that LC^NE^ neurons are involved in switching from sevoflurane anesthesia to wakefulness. In addition, using anterograde and retrograde tracing techniques, we confirmed that the mPFC receives projections from LC^NE^ neurons. We dug a bit deeper and discovered that firing up the LC-mPFC NE neurocircuit really packs a punch when it comes to boosting arousal during sevoflurane anesthesia.

## 2 Materials and methods

### 2.1 Animals

This study was approved by the Animal Ethics Committee of Zhejiang University and followed the National Institutes of Health’s guidelines for handling lab animals to a tee. Wild-type C57BL/6J mice were housed and bred in the SPF-level animal facility at the School of Medicine, Zhejiang University. To avoid any mix-ups from gender differences or the female mice’s estrous cycle, we stuck to using only male mice. And to keep everything on track, we made sure all the experiments happened between 9 a.m. and 4 p.m. That way, we had a tight rein on the conditions and could trust our results.

### 2.2 Sevoflurane anesthesia modeling and method of anesthesia induction and emergence time observation

C57BL/6J mice were placed in a well-sealed inhalation anesthesia chamber (dimensions: 20 cm in length, 10 cm in width, and 15 cm in height; model V101, RWD Life Sciences, Shenzhen, China). To familiarize the mice with the experimenter’s scent and the experimental environment, their cages were not changed 3–4 days before the experiment, and the mice were gently handled daily. On the day of the behavioral test, the mice were allowed to acclimate to the experimental environment for 30 minutes in advance. At both ends of the top of the anesthesia box, one end is connected to the gas input pipeline, with a gas flow rate of 2 liters per minute and an oxygen concentration of 60%, while the other end is connected to the gas recovery device. To maintain a comfortable and stable temperature within the anesthesia box during the experiment, a cotton pad was placed at the bottom of the box. Fifteen minutes prior to the start of the experiment, an electric heating pad was utilized to preheat the bottom of the box. This preheating step helped to create and sustain an optimal thermal environment for the mice throughout the experimental procedure. Immediately after the start of the experiment, the sevoflurane vaporizer was activated to maintain a sevoflurane vapor concentration of 2.5%. Additionally, the anesthesia box was rotated 90 degrees every 15 seconds. When the mouse could not right itself from an abnormal position to a normal four-limb stance, it indicated the loss of righting reflex (LORR), which was recorded as the anesthesia induction time. During the maintenance phase, corresponding experimental procedures were carried out in accordance with the experimental design. After 30 minutes of anesthesia, the sevoflurane vaporizer was shut down to terminate the inhalation anesthesia. To eliminate any residual sevoflurane from the anesthesia box and the associated pipelines, a gas mixture composed of 80% oxygen and 20% nitrogen was rapidly introduced at a steady flow rate of 2 liters per minute. When the mouse was able to voluntarily return to a normal four-limb stance from an abnormal position, it indicated the recovery of the righting reflex (RORR). The time elapsed from the cessation of anesthesia to the point of RORR was documented as the emergence time. To eliminate the influence of individual mouse differences and the experimental environment, the same batch of mice was used for both the control and experimental groups.

### 2.3 Pharmacological experiments

The selective norepinephrine (NE) reuptake inhibitor Atomoxetine was prepared using 0.9% normal saline. Forty mice from the same litter were randomly divided into five groups. Four groups of mice were intraperitoneally injected with different concentrations of Atomoxetine (5, 10, 15, or 20 mg/kg) and served as the experimental groups, while one group was injected with normal saline and served as the control group. One hour after the injection, the vaporizer was activated, and the mice were exposed to 2.5% sevoflurane for anesthesia induction and maintenance. Behavioral changes in the mice were observed, and the induction and awakening times were recorded. The groups were as follows: Vehicle group; Atomoxetine 5 mg/kg group; Atomoxetine 10 mg/kg group; Atomoxetine 15 mg/kg group; Atomoxetine 20 mg/kg group. The highly selective neurotoxin for the LC NE system, N-(2-chloroethyl)-N-ethyl-2-bromobenzylamine hydrochloride (DSP-4), was prepared using 0.9% normal saline. Forty mice from the same litter were randomly divided into five groups. Four groups were intraperitoneally injected with 50 mg/kg of DSP-4 and served as the experimental groups, while one group was injected with normal saline and served as the control group. Sevoflurane anesthesia experiments were conducted 1, 3, 5, or 10 days after DSP-4 injection to observe behavioral changes in the mice and record anesthesia induction and awakening times. The groups were as follows: Vehicle group; DSP-4 (1 d) group; DSP-4 (3 d) group; DSP-4 (5 d) group; DSP-4 (10 d) group.

### 2.4 Stereotaxic localization of mouse brain

After weighing and recording the body weight of the mice, C57BL/6J mice were anesthetized via intraperitoneal injection of 1% pentobarbital sodium (50 mg/kg). The mice were placed in a stereotactic frame (68018, RWD Life Science Co., Ltd., Shenzhen, China), and their skulls were leveled using ear bars and a nose clamp. During the surgery, a heating pad was used to maintain the body temperature of the anesthetized mice at 37°C. The eyes of the mice were coated with erythromycin ointment and then covered with sterile cotton balls to protect them from damage caused by intense light. Adjust the field of view and magnification of the microscope. Carefully trim the hair above the target brain region on the skull surface using tissue scissors. The area was disinfected from the inside out using a 75% alcohol swab. The cranial skin was then incised along the midline, and the mucosal tissue on the skull surface was removed. The sutures were wiped with 0.3% hydrogen peroxide to fully expose the anterior and posterior fontanelles of the mouse skull. Under the microscope, the injector tip was used for positioning. The tip was lifted and moved to the anterior fontanelle, where the coordinates were set to zero. The needle tip was then moved to the posterior fontanelle, and the height difference between the anterior and posterior fontanelles was adjusted to ensure it was no more than 0.03 millimeters. After leveling the anterior and posterior fontanelles, equidistant points (2.30 mm) on both sides of the anterior fontanelle were selected to determine the height. If the height difference was within 0.03 millimeters, it was considered that the skull had been leveled. Using the fourth edition of the mouse brain atlas (Paxinos and Franklin, 2013), the target brain regions were precisely located on the skull surface. For targeting the LC, the coordinates were: AP −5.41 mm, ML ±0.9 mm, DV −3.75 mm. For targeting the mPFC, the coordinates were: AP +1.90 mm, ML ±0.35 mm, DV −2.20 mm. After confirming the location, a micro handheld cranial drill was carefully used to create a hole with a diameter of 0.5-1 mm in the skull. The procedure requires great care. During drilling, it is crucial to avoid large blood vessels and brain parenchyma to prevent irreversible damage to brain tissue and to ensure the accuracy of the experimental results.

### 2.5 Viral microinjection into brain nuclei

Before viral injection, we filled the microsyringe with liquid paraffin to remove air bubbles and sealed the syringe-to-electrode connection with hot-melt glue, ensuring airtightness. We then leveled the skull and drilled a hole above the target brain region. Using a microinjection pump, we drew up a volume of viral solution slightly exceeding the experimental requirement. Under the microscope, we positioned the needle over the target area, ensuring the needle tip aligned with the drilled hole. We slowly inserted the needle to avoid bleeding, closely monitoring the mouse and keeping the needle on course. Once the needle reached the target, we held it in place for 5 minutes to allow the brain tissue to settle. The microinjection pump then delivered the viral solution at 40–50 nL/min, totaling 100 nL. During injection, we watched under the microscope to see the viral solution and liquid paraffin separate into layers and the interface descend. After injecting, we left the needle in place for 10 more minutes to allow the virus to diffuse. We then slowly withdrew the needle to prevent leakage into nearby brain regions. We cleaned the skull surface with a saline-soaked cotton ball, keeping the scalp clean and moist. Finally, we closed the scalp with absorbable sutures, applied erythromycin ointment to prevent infection, and placed the mouse on a temperature-controlled heating pad to recover. The mouse was returned to its cage only after it regained spontaneous activity.

### 2.6 Implantation of cannula in lateral ventricle and nucleus

We began by mounting the holder on the stereotactic frame, ensuring the fiber optic ferrule needle was firmly clamped. Under a stereomicroscope, we drilled three shallow holes (not penetrating the skull) at specific locations: to the left front of the anterior fontanelle and to the front of the left and right sides of the posterior fontanelle. Miniature screws were then secured into these holes, with about one-third of each screw protruding above the skull surface. Using the fiber optic tip for stereotactic positioning, we leveled the skull and drilled a hole directly above the target brain region. The holder was slowly lowered to insert the fiber optic, while carefully monitoring the mouse’s condition and ensuring no deviation occurred. The fiber optic was inserted to a depth of 0.1 mm above the viral injection site within the nucleus. Once the target position was reached, we gently cleaned the skull surface of bone debris and blood using a saline-soaked cotton ball to securely fix the fiber optic and prevent it from shifting. Dental cement was prepared by mixing dental resin with denture base liquid and was evenly applied around the fiber optic and over the skull surface. After the cement fully hardened, the holder was carefully removed to ensure the fiber optic had not shifted. Post-surgery, the mouse was placed on a temperature-controlled heating pad to recover and was returned to its cage once it regained spontaneous activity. After the experiment, we verified the positions of viral expression and fiber optic placement by checking viral fluorescence, ensuring the fiber optic was correctly positioned within the target brain region. Finally, we analyzed the behavioral data accordingly. This meticulous process ensured accurate placement and secure fixation of the fiber optic, crucial for reliable experimental outcomes.

### 2.7 Optogenetics

The blue light parameters in this experiment are as follows: wavelength of 465 nm, frequency of 20 Hz, and pulse duration of 20 ms. Fine-tune the laser’s output settings and measure the laser intensity at the jumper’s output end using a power meter. Maintain the laser intensity at approximately 15 mW. Blue light stimulation was administered from 10 minutes prior to the onset of sevoflurane administration until the Loss of Response to Reflexes (LORR), and again from 10 minutes before sevoflurane was discontinued until the Return of Response to Reflexes (RORR).

The virus rAAV2/9-DBH-CRE-WPRE-hGH polyA, which contains the DBH-specific promoter, was thoroughly mixed with the optogenetic virus rAAV2/9-hSyn-DIO-hChR2(H134R)-EGFP-WPRE-hGH polyA at a ratio of 1:2. Subsequently, the mixed viruses were injected into the bilateral LC regions of the mouse brain using a stereotactic apparatus (injection rate: 40-50 nL/min, injection volume: 100 nL). After allowing two weeks for viral infection to take effect, optical fiber cannulas were implanted. One week post-implantation, the laser was connected to the cannulas on the mouse’s head via an optical fiber patch cord, enabling the commencement of optogenetic experiments. The virus rAAV2/9-DBH-CRE-WPRE-hGH polyA, which contains the DBH-specific promoter, was thoroughly mixed with the optogenetic virus rAAV2/9-hSyn-DIO-hChR2(H134R)-EGFP-WPRE-hGH polyA at a ratio of 1:2. Subsequently, the mixed viruses were injected into the unilateral LC region of the mouse brain using a stereotactic apparatus (injection rate: 40-50 nL/min, injection volume: 100 nL). After allowing two weeks for the viral infection to establish, optical fiber cannulas were inserted into the contralateral downstream medial prefrontal cortex (mPFC) region. One week following the cannula implantation, the laser was connected to the cannulas on the mouse’s head via an optical fiber patch cord, thereby facilitating the initiation of optogenetic experiments.

### 2.8 Chemogenetics

The virus rAAV2/9-DBH-CRE-WPRE-hGH polyA, which contains the dopamine β-hydroxylase (DBH) specific promoter, was thoroughly mixed with the chemogenetic activation virus rAAV2/9-hSyn-DIO-hM3D(Gq)-mCherry-WPRE-hGH polyA, the chemogenetic inhibition virus rAAV2/9-hSyn-DIO-hM4D(Gi)-mCherry-WPRE-hGH polyA, or the non-functional element-containing control virus rAAV2/9-hSyn-DIO-mCherry-WPRE-hGH polyA at a ratio of 1:2. The mixed viruses were then injected into the bilateral LC regions of the mouse brain using a stereotactic apparatus (injection rate: 40-50 nL/min, injection volume: 100 nL). Three weeks after viral infection, chemogenetic experiments were conducted. One hour prior to the start of sevoflurane anesthesia, an intraperitoneal injection of CNO (1 mg/kg) was given to enable chemogenetic control of LC noradrenergic neurons. Using a stereotactic instrument, either the chemogenetic activation virus rAAV2/9-hSyn-DIO-hM3D(Gq)-mCherry-WPRE-hGH polyA, the chemogenetic inhibition virus rAAV2/9-hSyn-DIO-hM4D(Gi)-mCherry-WPRE-hGH polyA, or the non-functional control virus rAAV2/9-hSyn-DIO-mCherry-WPRE-hGH polyA was injected into the unilateral LC region of the mouse brain at a rate of 40–50 nL/min, with a total volume of 100 nL. Next, the retrograde virus rAAV2/Retro-DBH-CRE-WPRE-hGH polyA, equipped with the DBH-specific promoter, was co-injected with the virus rAAV2/9-hSyn-DIO-mCherry-WPRE-hGH polyA into the downstream mPFC region on the same side of the mouse brain. The injection was performed using a stereotactic apparatus at a rate of 40–50 nL/min, with a total volume of 100 nL. Viruses containing the DIO (Double-floxed inverse open reading frame) element only undergo recombination and express under the action of Cre recombinase. The retrograde virus rAAV2/Retro-DBH-CRE can retrogradely transport from axon terminals to the somas of upstream neurons. The DBH promoter drives the expression of Cre recombinase. DBH, an essential enzyme in neurotransmitter synthesis, catalyzes the conversion of dopamine to NE. Thus, the virus enables the expression of Cre recombinase specifically in the NE-ergic neurons that project from the LC to the mPFC, while the Cre recombinase, in turn, induces viral expression in the LC. Through the combination of these viral tools, we specifically labeled NE neurons in the LC that project to the mPFC and expressed chemogenetic viruses (hM3Dq or hM4Di) and a non-functional control virus (mCherry) in these neurons. Chemogenetic experiments were conducted three weeks after viral infection. Administering an intraperitoneal injection of CNO (1mg/kg) 1 hour before the initiation of sevoflurane anesthesia can achieve chemogenetic control of the LC-mPFC NEergic neuronal circuit.

### 2.9 Fiber photometry

Using a stereotactic apparatus, rAAV2/9-DBH-GCaMP6m-WPRE-hGH pA was injected into the LC region at a rate of 40–50 nL/min, with a total volume of 100 nL. Two weeks after viral infection, optical fiber cannulas were implanted. One week after the implantation of the optical fiber cannulas, fiber photometry experiments were conducted. Prior to the experiment, the output parameters of the laser were fine-tuned. Using a power meter, the laser intensity at the output end of the fiber patch cord was measured and adjusted to ensure that the 488 nm laser operated within a range of 20– 40 μW, and the 410 nm laser within 10–20 μW. Subsequently, the laser was connected to the cannulas on the mouse’s head via an optical fiber patch cord and interfaced with a computer recording software to capture calcium signal recordings.

In this experiment, the dual-color fiber photometry system employs dual-wavelength excitation light at 410 nm and 488 nm, capturing the resulting fluorescence signals simultaneously. The fluorescence signal changes elicited by the 410 nm excitation light reflect background noise and are used as a reference control. The fluorescence signals were segmented based on behavioral events and aligned with a time-zero point corresponding to specific behavioral events. The change in fluorescence intensity is characterized by the value of ΔF/F0 = (F - F0) / F0, where F represents the current fluorescence intensity and F0 represents the baseline fluorescence intensity. This metric is used to indicate the activity changes of LC noradrenergic neurons. The analysis results are represented using heat maps and peri-event plots.

### 2.10 Anterograde and Retrograde Tracing

Anterograde Tracing: HSV was injected into the LC region of the mouse brain using a stereotactic instrument at a rate of 40–50 nL/min, with a total volume of 100 nL. After allowing 2 days for expression, the mice were perfused to extract their brains. Retrograde Tracing: Using a stereotactic apparatus, CTB-555 was injected into the mPFC region of the mouse brain (injection rate: 40–50 nL/min, injection volume: 100 nL). After one week of expression, the mice were perfused to harvest their brains.

### 2.11 Immunohistochemistry

After the experiment, mice were given an intraperitoneal injection of 2% sodium pentobarbital (50 mg/kg) and positioned on a foam board. The thoracic cavity was opened to expose the heart. The perfusion needle was inserted into the left ventricle, and the right atrial appendage was cut to allow fluid outflow. First, phosphate-buffered saline (PBS) was used for perfusion until the liver turned pale. Then, 4% paraformaldehyde (PFA) was used for fixation until the limbs stopped convulsing and the body became rigid. The perfusion rate was carefully controlled to prevent fluid from entering the pulmonary circulation. Afterward, the brain was removed, immersed in 4% PFA fixative for 12–16 hours at 4°C, and then transferred to a 30% sucrose solution for 24 hours at 4°C for dehydration. The process was complete when the brain tissue sank.

After the perfusion, we decapitated the mouse and placed its brain in a 4% PFA fixative solution at 4°C for 12–16 hours. Then, the brain was transferred to a 30% sucrose solution at 4°C for 24 hours to dehydrate. Once the brain tissue sank, we knew it was ready for the next step. Next, we embedded the brain in OCT and cut it into 35–40 μm sections. These sections were placed in a 24-well plate and washed three times with PBS for 5 minutes each. After that, the sections were immersed in a blocking solution and incubated at room temperature for 2.5 hours. We then added the primary antibody dropwise and incubated the sections on a shaker in the refrigerator at 4°C overnight. The following day, the sections were washed three times in PBS for 10 minutes each. They were then placed in the secondary antibody solution and incubated for 1 hour at room temperature in the dark. After another three 15-minute PBS washes in a light-protected device, the sections were stained with DAPI for 7 minutes and washed again with PBS for 15 minutes to remove excess stain. Once dry, an antifade mounting medium was applied to seal the sections. Finally, we conducted fluorescence imaging using the VS120 virtual slide scanner system and the Nikon A1R confocal microscope. The immunofluorescent-positive cells and colocalized cells were quantified and analyzed using ImageJ and NIS-Elements Viewer. This meticulous process ensures that the brain tissue is properly prepared and analyzed, providing valuable insights for further research.

### 2.12 GABAA-R knockdown

We microinjected 80 nL of rAAV-Dbh-EGFP-S’miR-30a-shRNA (GABAA receptor)–3’-miR30a-WPRES (viral titer: 5E+12 vg/mL, Brain VTA Technology Co, Ltd) into the LC. Four weeks later, the mice were euthanized and perfused. This was done to count the GABAA-R positive and tyrosine hydroxylase-positive (TH+) cells, which helped us see how well the GABAA-R knockdown worked.

### 2.13 Quantification and statistical analysis

The experimental data were analyzed using GraphPad Prism 9.0 and SPSS version 26.0 statistical analysis software. Prior to analysis, all experimental data were subjected to the Shapiro-Wilk normality test and Levene’s test for homogeneity of variances. For comparisons between two groups: If the data followed a normal distribution and had homogeneous variances, an independent samples t-test (Student’s t-test) was used. If the data followed a normal distribution but had heterogeneous variances, Welch’s t-test was used. If the data did not follow a normal distribution, the Mann-Whitney U test was used. For comparisons involving three or more groups: If the data followed a normal distribution and had homogeneous variances, one-way analysis of variance (One-way ANOVA) was used. If the data followed a normal distribution but had heterogeneous variances, Welch’s ANOVA was used. If the data did not follow a normal distribution, the Kruskal-Walli’s test was used. When conducting multiple comparisons, post hoc tests are required. Experimental data are expressed as mean ± standard error of the mean (Mean ± SEM). A P < 0.05 is considered statistically significant.

## 3 Results

### 3.1 Pharmacological modulation of central NE levels affects sevoflurane anesthesia-to-arousal transition

To assess LC^NE^ neurons involvement in sevoflurane anesthesia, we analyzed c-Fos expression in these neurons. Fluorescence images revealed a significant reduction in the number of c-Fos-positive cells in the sevoflurane group compared to the control group (P<0.0001, Fig. 1A-B), suggesting anesthesia-induced suppression. This result indicated that LC^NE^ neurons may regulate sevoflurane anesthesia-induced consciousness transitions. To investigate the role of central NE levels in the sevoflurane anesthesia-to-arousal transition process, we conducted pharmacological intervention experiments (Fig. 1C). We found that IP injection of atomoxetine (20 mg/kg), a central selective NE reuptake inhibitor, significantly increased the number of c-Fos (+)/TH (+) cells in the LC (P<0.0001, Fig. 1D-E), prolonged the anesthesia induction time (P<0.001, Fig. 1F) and shortened the arousal time after sevoflurane administration (P<0.0001, Fig. 1G). Conversely, IP injection of 50 mg/kg N-(2-chloroethyl)-N-ethyl-2-bromobenzylamine hydrochloride (DSP-4) (Fig. 1H), a highly selective NEergic neurotoxin, decreased the number of TH+ cells in the LC, with the most significant reduction observed on day 5 or day 10 post-injection (P<0.0001, Fig. 1I-J). This DSP-4 injection also significantly shortened the anesthesia induction time (P<0.001, P<0.01, Fig. 1K) and prolonged the arousal time after sevoflurane administration (P<0.0001, Fig. 1L). These results collectively suggest that the LC NE system plays a pivotal role in modulating the alterations in consciousness induced by sevoflurane.

**Figure 1.**
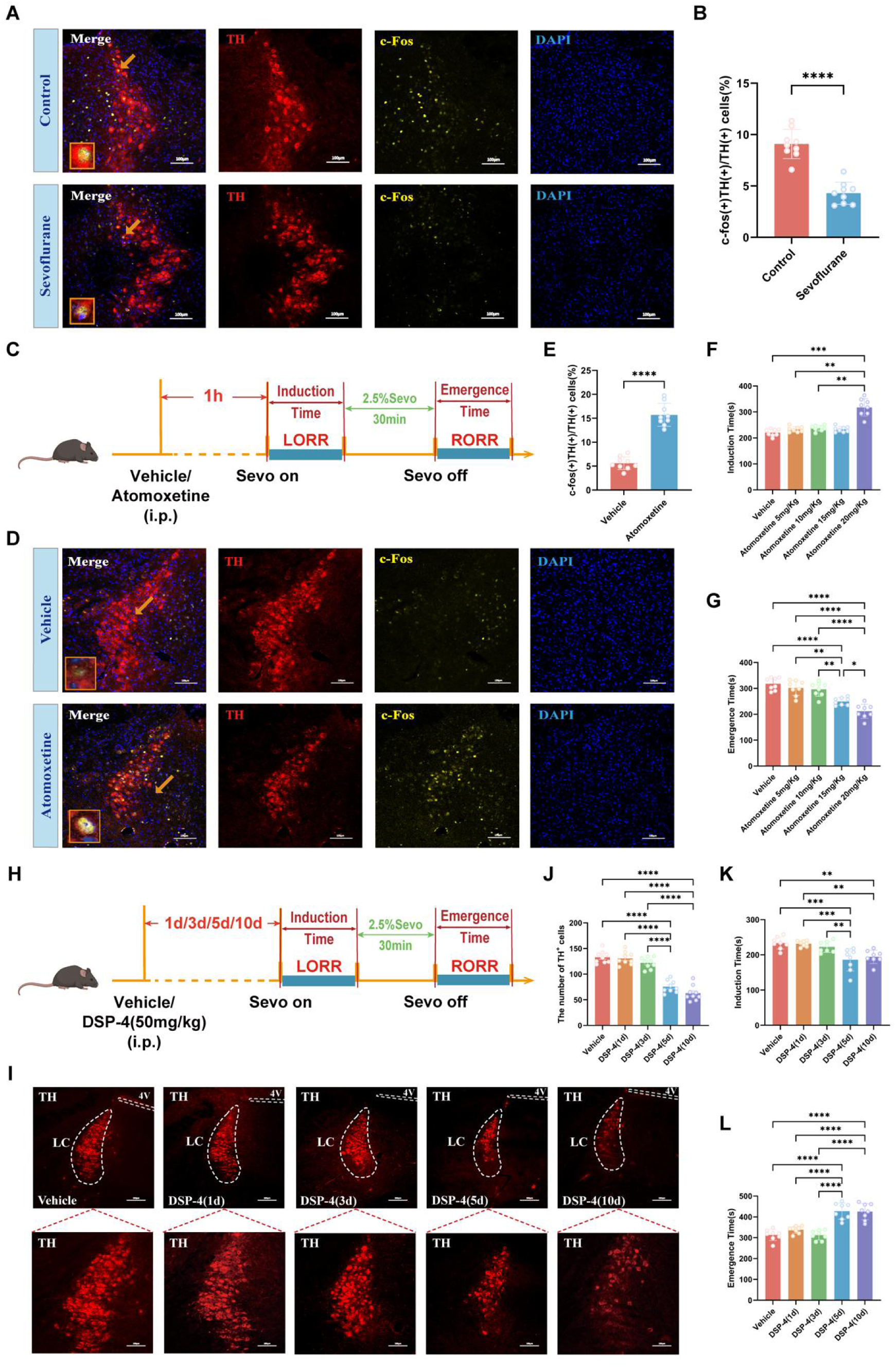
Preliminary pharmacological studies have explored LC^NE^ neurons promote emergence from sevoflurane anesthesia. **(A)** Representative coronal brain slice of LC, showing the staining for c-Fos, TH and DAPI with or without sevoflurane. **(B)** The quantification of c-fos(+) TH(+)/TH(+) cells in the LC with or without sevoflurane. **(C)** Experimental protocol for IP injection of different doses of Atomoxetine. **(D)** Representative coronal brain slice of LC, showing the staining for c-Fos, TH and DAPI with or without IP injection of Atomoxetine. **(E)** The quantification of c-fos(+) TH(+)/TH(+) cells in the LC with or without IP injection of Atomoxetine. **(F, G)** Induction time and emergence time for IP injection in Vehicle, 5 mg/kg Atomoxetine, 10 mg/kg Atomoxetine, 15 mg/kg Atomoxetine and 20 mg/kg Atomoxetine groups. **(H)** Experimental protocol for IP injection of 50 mg/kg DSP-4. **(I)** Representative coronal brain slice of LC, showing the staining for TH in Vehicle, DSP-4(1d), DSP-4(3d), DSP-4(5d) and DSP-4(10d) groups. **(J)** The quantification of the numbers of TH(+) cells in the LC in Vehicle, DSP-4(1d), DSP-4(3d), DSP-4(5d) and DSP-4(10d) groups. **(K, L)** Induction time and emergence time for IP injection in Vehicle, DSP-4(1d), DSP-4(3d), DSP-4(5d) and DSP-4(10d) groups. ****p<0.0001; ***p<0.001; **p<0.01; *p<0.05; i.p., Intraperitoneal injection; Sevo, Sevoflurane; LORR, Loss of Righting Reflex; RORR, Recovery of Righting Reflex

### 3.2 LC^NE^ neurons contribute to regulating consciousness transition from sevoflurane anesthesia

To further quantify the neuronal activity of LC^NE^ neurons at different states during sevoflurane anesthesia, we performed calcium imaging. We injected AAV2/9-DBH-GCaMP6m-WPRE-hGH pA with a specific promoter into the bilateral LC of C57BL/6J mice to express the GCaMP6m calcium indicator specifically in NEergic neurons and examined changes in GCaMP6m fluorescence signal through an optical fiber located in the LC (Fig. 2A-B, E). confirming sufficient expression of the calcium virus in LC^NE^ neurons, specifically localized to the LC region (Fig. 2C-D). During the LORR to RORR phase, we found that the ΔF/F peak of calcium signaling in LC^NE^ neurons was significantly reduced (P<0.01, P<0.05, Fig. 2F-H) compared with other phases, and then increased after RORR (P<0.001, Fig. 2F-H).These findings demonstrated LC^NE^ neuronal inhibition during sevoflurane anesthesia and gradual reactivation post-cessation, supporting their role in anesthesia-arousal regulation.

**Figure 2.**
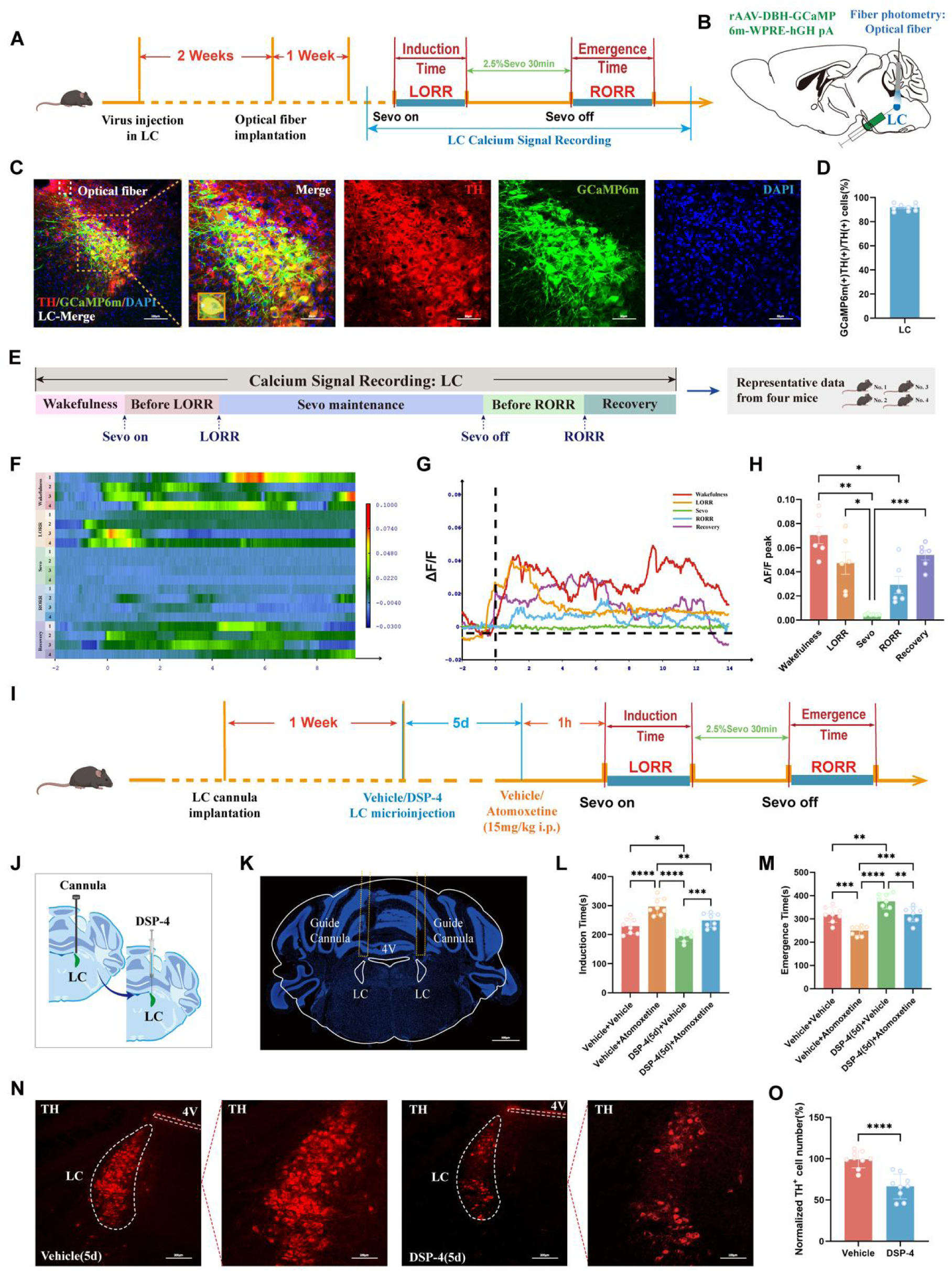
Fiber photometry and microinjection intra-LC specifically revealed the pro-arousal effects of LC^NE^ neurons. **(A)** Experimental protocol for calcium signal recording of LC^NE^ neurons. **(B)** Schematic of the position of virus injection and optic fibers implantation in the LC. **(C)** Representative coronal brain slice, showing the location of virus injection, and representative images of staining for DAPI, TH and GCaMP6m in the LC. **(D)** The quantification of GCaMP6m(+) TH(+)/TH(+) cells in the LC. **(E)** Schematic diagram of the different stages of the calcium signal recording in the LC and the sources of statistical data. **(F-H)** Heatmap and statistical diagram show the activity wave of calcium signals and the peak ΔF/F in the LC during the Wakefulness, Before LORR, Sevoflurane, Before RORR and Recovery five stages. **(I)** Experimental protocol for intra-LC microinjection of DSP-4 and/or Atomoxetine. **(J)** Schematic of the position of the cannula implantation and microinjection of DSP-4 in the LC. **(K)** Representative image of the track of cannula implanted into the LC. **(L, M)** Induction time and emergence time for microinjection in Vehicle + Vehicle, Vehicle + Atomoxetine, DSP-4(5d) + Vehicle and DSP-4(5d) + Atomoxetine groups. **(N)** Representative coronal brain slice of LC, showing the staining for TH in Vehicle(5d) and DSP-4(5d) groups. **(O)** The quantification of the numbers of TH(+) cells in the LC in Vehicle(5d) and DSP-4(5d) groups. ****p<0.0001; ***p<0.001; **p<0.01; *p<0.05; Sevo, Sevoflurane; LORR, Loss of Righting Reflex; RORR, Recovery of Righting Reflex

To delve deeper into LC^NE^ neurons in recovery after sevoflurane anesthesia, we artificially intervened in LC^NE^ neurons (Fig. 2I). After the anesthesia experimental protocol, we performed immunofluorescence staining and confirmed that the cannulas implantation position was accurate (Fig. 2J-K). Furthermore, microinjecting DSP-4 into LC not only significantly extended the emergence time from sevoflurane-induced loss of consciousness but also attenuated the arousal-promoting effect of intraperitoneally administered atomoxetine (P<0.01, P<0.001, Fig. 2L, M). We microinjected DSP-4 into LC to specifically degrade LC^NE^ neurons and found a significant decrease in the number of TH+ neurons in the LC of mice 5 days after the DSP-4 microinjection (P<0.0001, Fig. 2N, O). These suggest that intranuclear administration of DSP-4 to specifically ablate LCNE neurons postpones recovery from sevoflurane anesthesia.

### 3.3 Specific regulation of LC^NE^ neurons activity affects the sevoflurane anesthesia-arousal consciousness transition process

The aforementioned experiments preliminarily indicated that LC^NE^ neurons might be involved in regulating the loss of consciousness induced by sevoflurane anesthesia and the arousal process. To further explore the role of LC^NE^ neurons in the sevoflurane anesthesia-arousal process, we employed chemogenetics to specifically regulate the activity of LC^NE^ neurons and observe changes in the induction and emergence time of sevoflurane anesthesia. Firstly, we used chemogenetics to specifically activate LC^NE^ neurons. We injected a mixture of rAAV2/9-DBH-CRE and rAAV2/9-hSyn-DIO-hM3D(Gq)-mCherry viruses (hM3Dq) or rAAV 2/9-DBH-CRE and rAAV2/9-hSyn-DIO-mCherry viruses (mCherry) into bilateral LC brain regions (Fig. 3A-B). The results showed that both the chemogenetic activation virus hM3D(Gq)-mCherry and the control virus mCherry had a high co-expression rate with TH (Fig. 3C-D), and overall, the viruses were fully expressed in LC^NE^ neurons and confined to the LC. We found that immunofluorescence images revealed a significant increase in c-Fos expression in LC^NE^ neurons after chemogenetic activation (P<0. 0001, Fig. 3E-F). Moreover, after activating the bilateral LC brain regions, IP injection of 1 mg/kg CNO significantly prolonged the induction time of sevoflurane anesthesia (P<0.01, Fig. 3G) and markedly shortened the arousal time (P<0.001, Fig. 3H).

**Figure 3.**
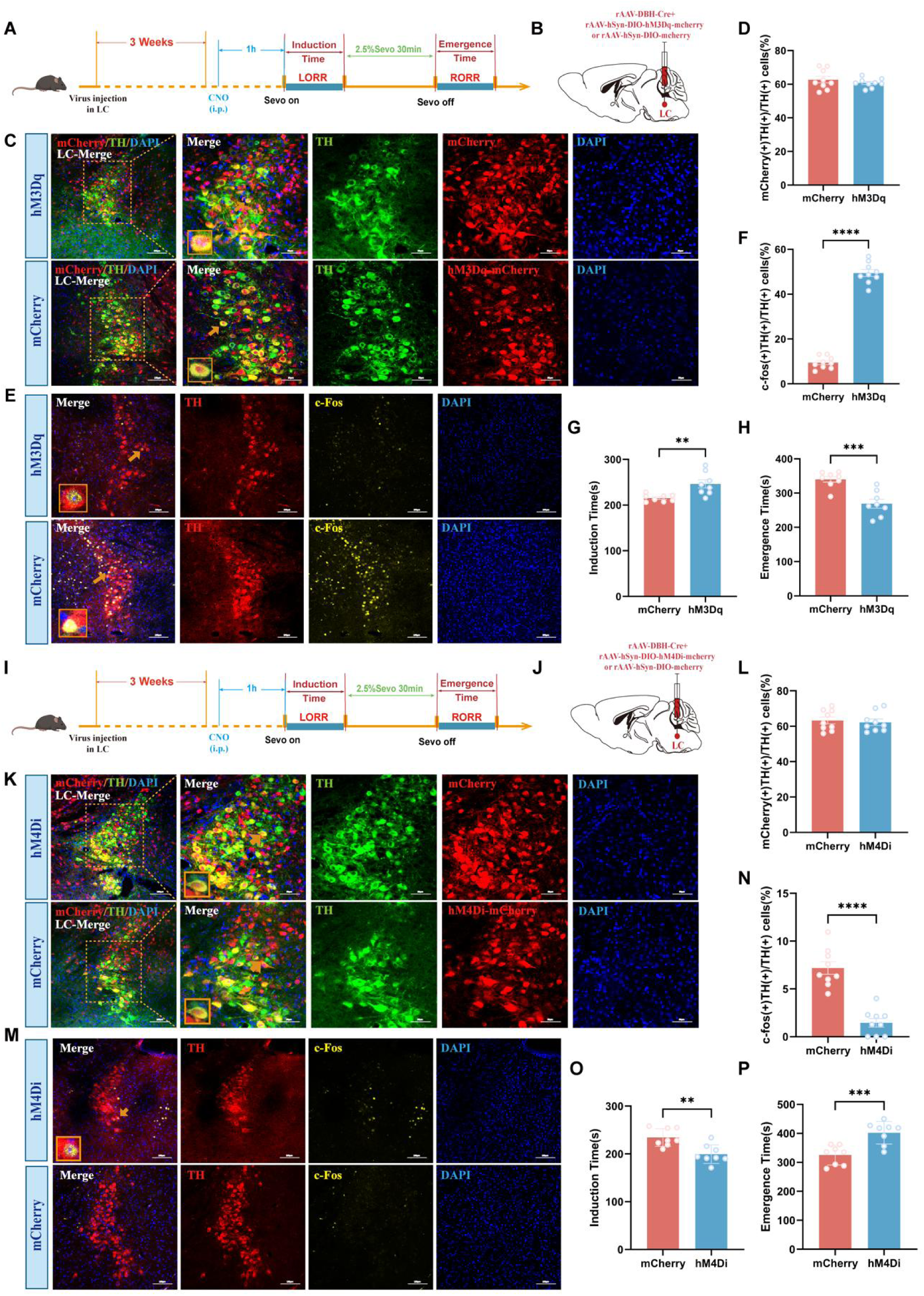
Chemogenetic activation and inhibition of LC^NE^ neurons promote arousal from sevoflurane anesthesia. **(A)** Experimental protocol for chemogenetic activation of LC^NE^ neurons. **(B)** Schematic of the position of the virus injection in the LC. **(C)** Representative images show the location of virus injection in the LC and the co-expression of hM3Dq-mCherry/mCherry, TH and DAPI in mCherry and hM3Dq groups. **(D)** The quantification of mCherry(+)TH(+)/TH(+) cells in the LC in mCherry and hM3Dq groups. **(E)** Representative images show the co-expression of and staining for c-Fos, TH and DAPI in the LC in mCherry and hM3Dq groups. **(F)** The quantification of c-fos(+)TH(+)/TH(+) cells in the LC in mCherry and hM3Dq groups. **(G, H)** Induction time and emergence time for chemogenetic activation in mCherry and hM3Dq groups. **(I)** Experimental protocol for chemogenetic inhibition of LC^NE^ neurons. **(J)** Schematic of the position of the virus injection in the LC. **(K)** Representative images show the location of virus injection in the LC and the co-expression of hM4Di-mCherry /mCherry, TH and DAPI in mCherry and hM4Di groups. **(L)** The quantification of mCherry(+)TH(+)/TH(+) cells in the LC in mCherry and hM4Di groups. **(M)** Representative images show the co-expression of and staining for c-Fos, TH and DAPI in the LC in mCherry and hM4Di groups. **(N)** The quantification of c-fos(+) TH(+)/TH(+) cells in the LC in mCherry and hM4Di groups. **(O, P)** Induction time and emergence time for chemogenetic inhibition in mCherry and hM4Di groups. ****p<0.0001, ***p<0.001, **p<0.01; i.p., Intraperitoneal injection; Sevo, Sevoflurane; LORR, Loss of Righting Reflex; RORR, Recovery of Righting Reflex

Concurrently, we conducted chemogenetic inhibition experiments. We injected a mixture of rAAV2/9-DBH-CRE and rAAV2/9-hSyn-DIO-hM4D(Gi)-mCherry viruses (hM4Di) or rAAV2/9-DBH-CRE and rAAV2/9-hSyn-DIO-mCherry viruses (mCherry) into bilateral LC brain regions (Fig. 3I-J). The results showed that both the chemogenetic inhibited virus hM4D(Gi)-mCherry and the control virus mCherry had a high co-expression rate with TH (Fig. 3K-L), and overall, the viruses were fully expressed in LC^NE^ neurons and confined to the LC. The results showed that chemogenetic inhibition significantly reduced c-Fos expression in LC^NE^ neurons (P<0.0001, Fig. 3M-N). Additionally, after chemically inhibiting the bilateral LC brain regions, IP injection of 1 mg/kg CNO significantly shortened the induction time of sevoflurane anesthesia (P<0.01, Fig. 3O) and markedly prolonged the arousal time (P<0.001, Fig. 3P). These findings further underscore the crucial role of these neurons in the sevoflurane anesthesia-arousal consciousness transition process.

### 3.4 The existence of anatomical neural projections from LC^NE^ neurons to the mPFC

In the experiments mentioned above, we initially confirmed the involvement of LCNE neurons in regulating the sevoflurane anesthesia-to-arousal process. The next question arises: how do LC^NE^ neurons exert this effect, and what are their downstream projection pathways? LC^NE^ neurons project extensively to the entire brain, including the mPFC, a crucial region involved in the regulation of cognitive functions, arousal states, and consciousness alterations. Therefore, we hypothesize that the mPFC may serve as a downstream brain area targeted by the LC in regulating anesthesia-to-wakefulness consciousness transitions, and that the LC-mPFC neural circuit plays a significant role in this process. Preliminarily, we hypothesize that the LC-mPFC NE neural circuit participates in regulating the sevoflurane anesthesia-to-arousal transition. Our experimental results show that chemogenetic activation of LC^NE^ neurons promotes arousal from sevoflurane anesthesia (Fig. 4A). Concurrently, immunofluorescence staining of the mPFC brain region revealed a significant increase in c-Fos expression (P<0.0001, Fig. 4B), indicating marked activation of the mPFC. This finding suggests a functional connection between LC^NE^ neurons and the mPFC.

**Figure 4.**
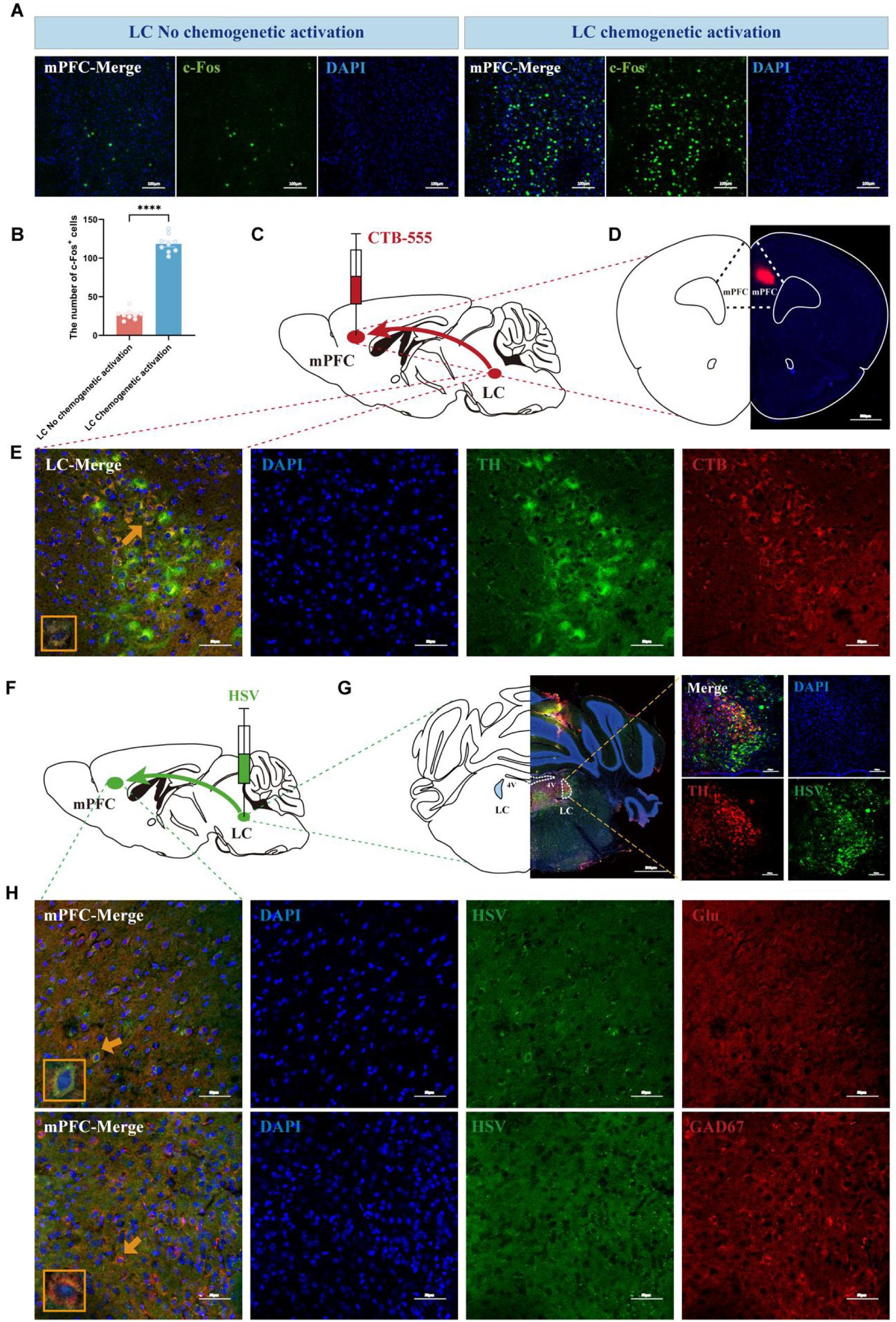
The existence of NEergic projection relationship between the LC and mPFC. **(A)** Representative coronal brain slice of mPFC, showing the staining for c-Fos and DAPI with or without LC chemogenetic activation. **(B)** The quantification of the numbers of c-Fos(+) cells in the mPFC with or without LC chemogenetic activation. **(C)** Schematic diagram of the injection position of retrograde labeling CTB-555. **(D)** Representative coronal brain slice of retrograde labeling CTB-555 in the mPFC. **(E)** Representative images show the co-expression of and staining for CTB, TH and DAPI in the LC. **(F)** Schematic diagram of the anterograde tracing by injecting anterograde tracing virus HSV into LC. **(G)** Representative coronal brain slice, showing the location of HSV injection and the staining of HSV, TH and DAPI in the LC. **(H)** Representative images show the co-expression and staining of HSV, Glu/GAD67 and DAPI in the mPFC. ****p<0.0001

To clarify the anatomical neural projection relationship between the LC and mPFC, we first injected HSV virus into the LC for anterograde tracing to observe whether neurons in the mPFC were labeled (Fig. 4C). We identified the HSV-infected region within the LC (Fig. 4D) and observed green, fluorescent signals from upstream LC HSV in the mPFC (Fig. 4E). We further confirmed the projection of fibers from LC^NE^ neurons to both glutamatergic and GABAergic neurons in the mPFC by immunofluorescence co-staining of HSV+ neurons in the mPFC with GAD67 or Glu (Fig. 15E). To further validate the neural projection relationship between the LC and mPFC, we injected CTB-555 virus into the mPFC for retrograde tracing (Fig. 4F). We confirmed the virus-infected region within the mPFC (Fig. 4G) and observed red fluorescent signals from downstream mPFC CTB in the LC (Fig. 4H). Subsequent immunofluorescence co-staining of CTB+ neurons in the mPFC with TH further confirmed the projection of fibers from LC^NE^ neurons to the mPFC (Fig. 4H). In summary, we have doubly verified the presence of anatomical neural projections from LC^NE^ neurons to the mPFC using both anterograde and retrograde tracing techniques.

### 3.5 Optogenetic activation of LC-mPFC neural circuitry extends sevoflurane anesthesia induction time and shortens awakening time

To precisely manipulate LC^NE^ neurons, we utilized optogenetics to activate them. We injected a mixture of rAAV2/9-DBH-CRE and rAAV2/9-hSyn-DIO-hChR2(H134R)-EGFP viruses into bilateral LC brain regions to specifically infect NE neurons within the LC, and optical fibers were implanted at the injection sites (Fig. 5A-B). In this experiment, blue light stimulation was applied from 10 minutes before sevoflurane administration to LORR and from 10 minutes before sevoflurane cessation to RORR, with a laser intensity of approximately 15 mW. The results showed that the optogenetic virus hChR2(H134R)-EGFP had a high co-expression rate with TH (Fig. 5C, E), and in general the viruses were fully expressed in LC^NE^ neurons and was confined to the LC. The fluorescent images revealed a significant increase in c-Fos expression in LC^NE^ neurons after light stimulation (P<0.0001, Fig. 5D, F). Our results demonstrated that optogenetic activation of bilateral LC brain regions significantly prolonged the induction time of sevoflurane anesthesia in mice (P<0.01, Fig. 5G) and markedly shortened the arousal time (P<0.0001, Fig. 5H).

**Figure 5.**
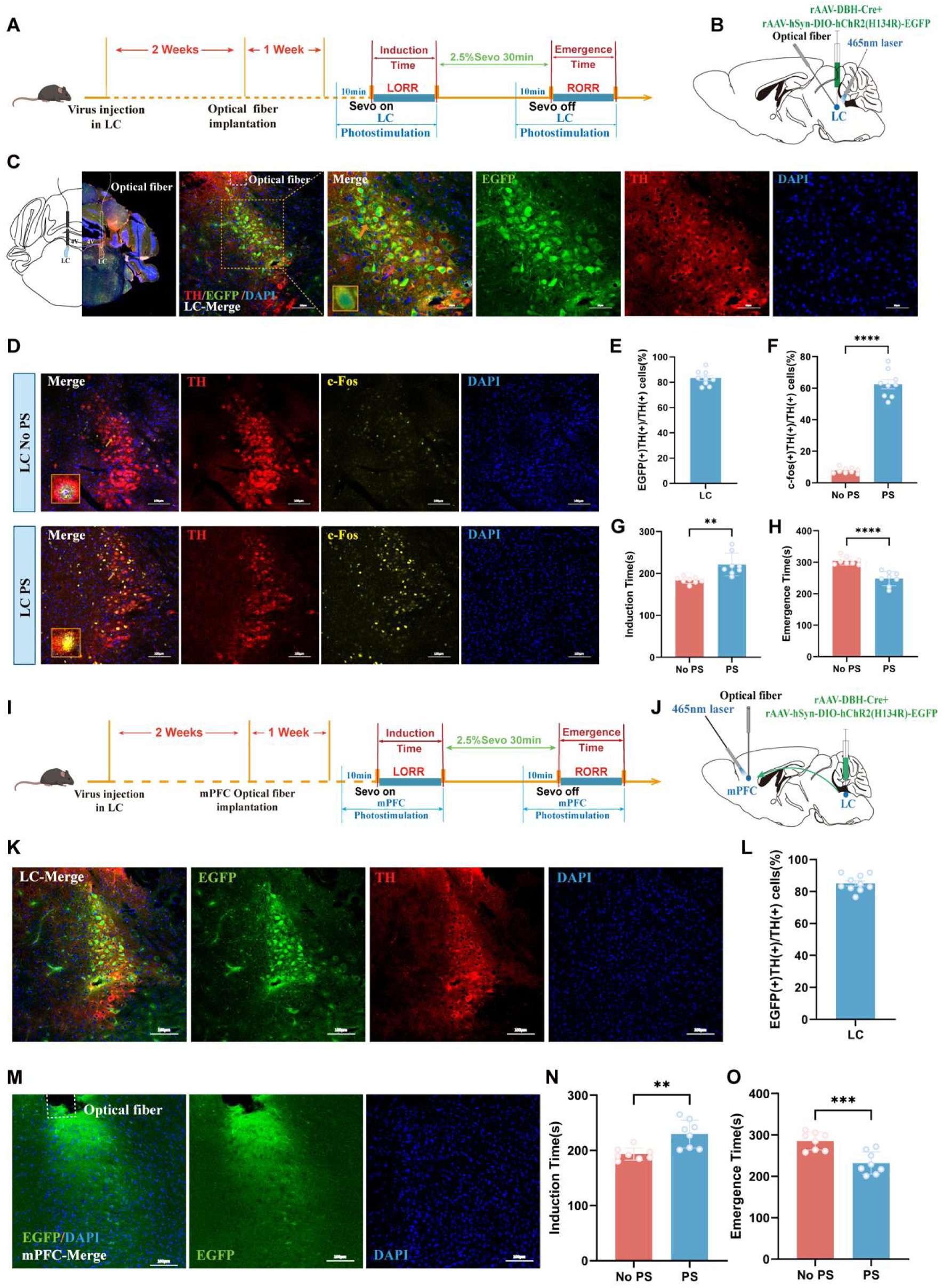
Optogenetic activation of the LC-mPFC NEergic neural circuit promotes arousal from sevoflurane anesthesia. **(A)** Experimental protocol for optogenetic activation of LC^NE^ neurons. **(B)** Schematic of the position of the virus injection and optic fibers implantation in the LC. **(C)** Representative coronal brain slice, showing the location of virus injection and the staining of EGFP, TH and DAPI in the LC. **(D)** Representative images of staining for c-Fos, TH and DAPI in the LC with and without LC photostimulation. **(E)** The quantification of EGFP(+)TH(+)/TH(+) cells in the LC. **(F)** The quantification of c-fos(+)TH(+)/TH(+) cells in the LC with and without LC photostimulation. **(G, H)** Induction time and emergence time for optogenetic activation with and without LC photostimulation. **(I)** Experimental protocol for long-range optogenetic activation of the LC-mPFC neural circuit. **(J)** Schematic of the position of the virus injection in the LC and optic fibers implantation in the mPFC. **(K)** Representative images, showing the location of virus injection and the staining of EGFP, TH and DAPI in the LC. **(L)** The quantification of EGFP(+) TH(+)/TH(+) cells in the LC. **(M)** Representative images of staining for EGFP and DAPI in the mPFC **(N, O)** Induction time and emergence time for optogenetic activation with and without mPFC photostimulation. ****p<0.0001; ***p<0.001; **p<0.01; Sevo, Sevoflurane; LORR, Loss of Righting Reflex; RORR, Recovery of Righting Reflex; PS, Photostimulation

To more precisely regulate the LC-mPFC neural circuitry, we used optogenetics to activate LC^NE^ terminals in the mPFC. A mixture of rAAV2/9-DBH-CRE and rAAV2/9-hSyn-DIO-hChR2(H134R)-EGFP viruses was injected into the unilateral LC, followed by the implantation of optical fibers in the ipsilateral downstream mPFC brain region (Fig. 5I-J). In this experiment, blue light stimulation was applied from 10 minutes before sevoflurane administration to LORR and from 10 minutes before sevoflurane cessation to RORR, with a laser intensity of approximately 15 mW. The results showed a high co-expression rate of the optogenetic virus hChR2(H134R)-EGFP with TH (Fig. 5K-M). Additionally, when blue light was used to activate the NE terminals of the LC-mPFC neural circuitry, the sevoflurane anesthesia induction time was significantly prolonged (P<0.01, Fig. 5N), and the awakening time was markedly shortened (P<0.001, Fig. 5O) compared to the non-light group. These findings further demonstrate the involvement of the LC-mPFC neural circuitry in regulating the consciousness transitions during sevoflurane anesthesia-to-awakening and the pro-awakening effect of activating this neural circuitry.

### 3.6 The LC-mPFC neural circuit involves in regulating the transition of consciousness during sevoflurane anesthesia and awakening

To further determine whether the neural projections between the LC and mPFC are related to sevoflurane anesthesia-arousal, we first used chemogenetics to activate the LC-mPFC neural circuit. We injected chemogenetic activation virus rAAV2/9-hSyn-DIO-hM3D(Gq)-mCherry (hM3Dq) or control virus rAAV2/9-hSyn-DIO-mCherry (mCherry) into the unilateral LC brain region, and co-injected retrograde virus rAAV2/Retro-TH-Cre with Cre-dependent virus rAAV2/9-hSyn-DIO-mCherry into the ipsilateral downstream mPFC brain region (Fig. 6A-B). We confirmed the infection of the retrograde virus in the mPFC (Fig. 6C) and found some neurons co-labeled with mCherry and TH in the LC of both groups of mice (Fig. 6D-E). Additionally, we observed that IP injection of 1 mg/kg CNO after chemogenetic activation of the unilateral LC brain region significantly prolonged the sevoflurane anesthesia induction time (P<0.01, Fig. 6F) and shortened the arousal time (P<0.001, Fig. 6G).

**Figure 6.**
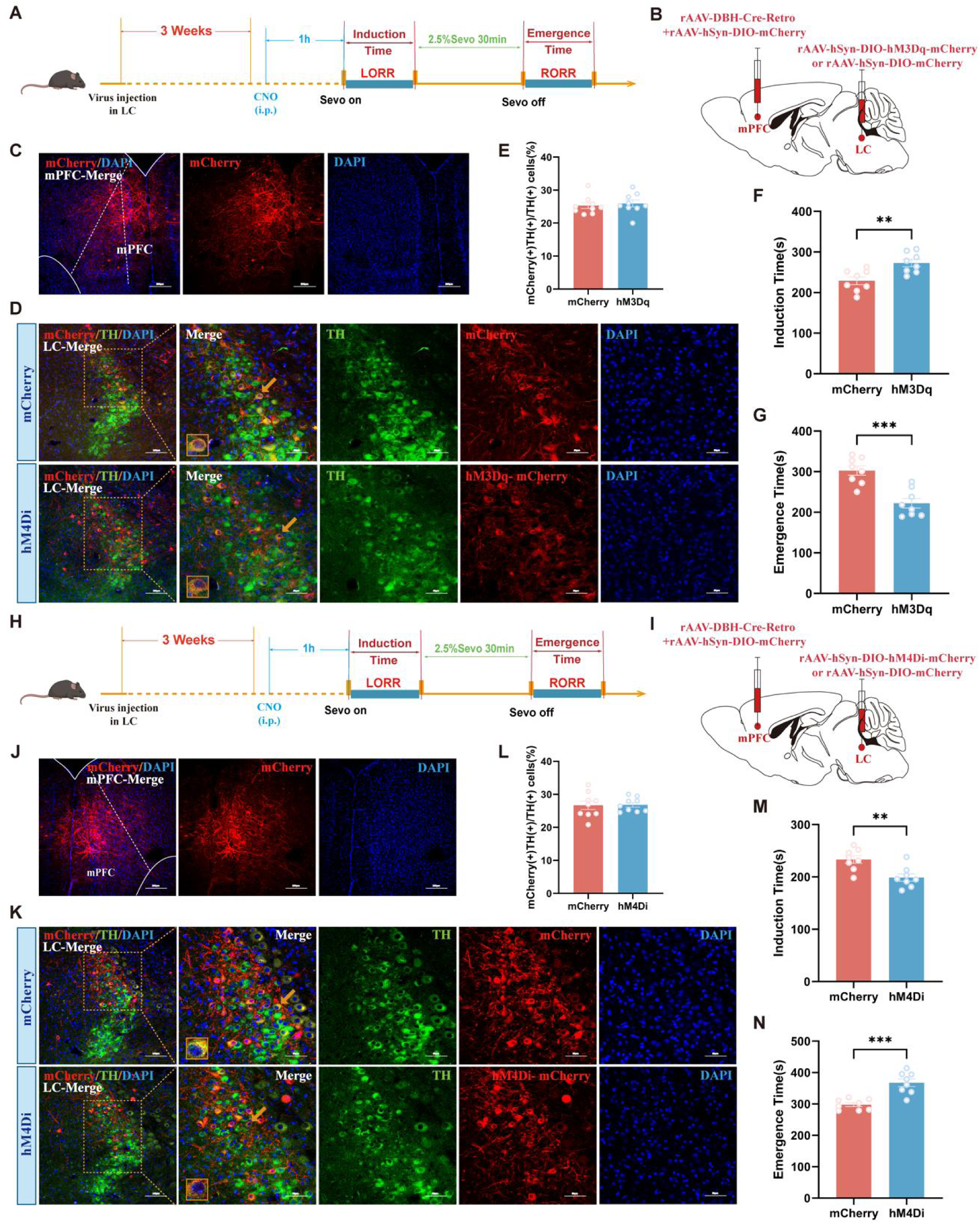
Chemogenetic activation of the LC-mPFC NEergic neural circuit promotes arousal from sevoflurane anesthesia. **(A)** Experimental protocol for long-range chemogenetic activation of the LC-mPFC neural circuit. **(B)** Schematic of the position of the virus injection in the LC and mPFC. **(C)** Representative images show the location of retro virus injection in the mPFC and the co-expression of mCherry and DAPI. **(D)** Representative images show the location of virus injection in the LC and the co-expression of hM3Dq-mCherry/mCherry, TH and DAPI in mCherry and hM3Dq groups. **(E)** The quantification of mCherry(+)TH(+)/TH(+) cells in the LC in mCherry and hM3Dq groups. **(F, G)** Induction time and emergence time for chemogenetic activation in mCherry and hM3Dq groups. **(H)** Experimental protocol for chemogenetic inhibition of the LC-mPFC neural circuit. **(I)** Schematic of the position of the virus injection in the LC and mPFC. **(J)** Representative images show the location of retro virus injection in the mPFC and the co-expression of mCherry and DAPI. **(K)** Representative images show the location of virus injection in the LC and the co-expression of hM4Di-mCherry/mCherry, TH and DAPI in mCherry and hM4Di groups. **(L)** The quantification of mCherry(+)TH(+)/TH(+) cells in the LC in mCherry and hM4Di groups. **(M, N)** Induction time and emergence time for chemogenetic inhibition in mCherry and hM4Di groups. ***p<0.001, **p<0.01, i.p., Intraperitoneal injection; Sevo, Sevoflurane; LORR, Loss of Righting Reflex; RORR, Recovery of Righting Reflex

To further confirm the involvement of LC-mPFC neural projections in sevoflurane anesthesia-to-awakening transitions, we conducted chemogenetic inhibition experiments. The chemogenetic inhibition virus rAAV2/9-hSyn-DIO-hM4D(Gi)-mCherry (hM4Di) or the control virus rAAV2/9-hSyn-DIO-mCherry (mCherry) was injected into the unilateral LC brain region. Concurrently, the retrograde virus rAAV2/Retro-TH-Cre and the Cre-dependent virus rAAV2/9-hSyn-DIO-mCherry were co-injected into the ipsilateral downstream mPFC brain region (Fig. 6H-I).We confirmed the infection of the retrograde virus in the mPFC brain region (Fig. 6J) and observed some mCherry and TH co-labeled neurons in the LC of both groups of mice (Fig. 6K-L). Furthermore, we found that IP injection of 1 mg/kg CNO significantly shortened the sevoflurane anesthesia induction time (P<0.01, Fig. 6M) and prolonged the awakening time (P<0.001, Fig. 6N) after chemogenetic inhibiting the unilateral LC brain region. Together, these results indicate the crucial role of the LC-mPFC neural circuitry in the transitions of consciousness during sevoflurane anesthesia-to-awakening.

### 3.7 α1-AR mediates the regulation of LC-mPFC neural circuitry in sevoflurane anesthesia-to-awakening consciousness transitions

α1-AR is widely distributed in the central nervous system, including the mPFC brain region. To determine the role of α1-AR in the LC-mPFC neural circuitry during sevoflurane anesthesia-to-awakening consciousness transitions, we employed pharmacological methods (Fig. 7A). After behavioral testing, we performed immunofluorescence staining and confirmed that the cannulas implantation position was accurate (Fig. 7B-C). Microinjection of Prazosin into the mPFC nucleus to antagonize α1-AR revealed that, compared to the control group, microinjection of α1-AR antagonist Prazosin into the mPFC nucleus shortened sevoflurane anesthesia induction time (P<0.05, Fig. 7D) and prolonged anesthesia awakening time (P<0.01, Fig. 7E).To further verify whether α1-AR participates in the LC-mPFC neural circuitry regulation of sevoflurane anesthesia-to-awakening consciousness transitions, we injected a mixture of rAAV2/9-DBH-CRE and rAAV2/9-hSyn-DIO-hM3D(Gq)-mCherry viruses (hM3Dq) or rAAV2/9-DBH-CRE and rAAV2/9-hSyn-DIO-mCherry viruses (mCherry) into the bilateral LC brain regions and implanted cannulas in the mPFC brain region (Fig. 7H-I). The results showed that chemogenetic activating LC^NE^ neurons significantly extended sevoflurane anesthesia induction time and promoted anesthesia awakening time (P<0.01, P<0.001, Fig. 7J-K), whereas microinjection of α1AR antagonist Prazosin into the mPFC significantly shortened sevoflurane anesthesia induction time and inhibited anesthesia awakening (P<0.05, P<0.05, Fig.7J-K). Notably, blocking α1AR with Prazosin significantly reversed the effects of chemogenetic activating LC^NE^ neurons on sevoflurane anesthesia induction time extension and anesthesia awakening time shortening (P<0.0001, P<0.0001, Fig. 7J-K).

**Figure 7.**
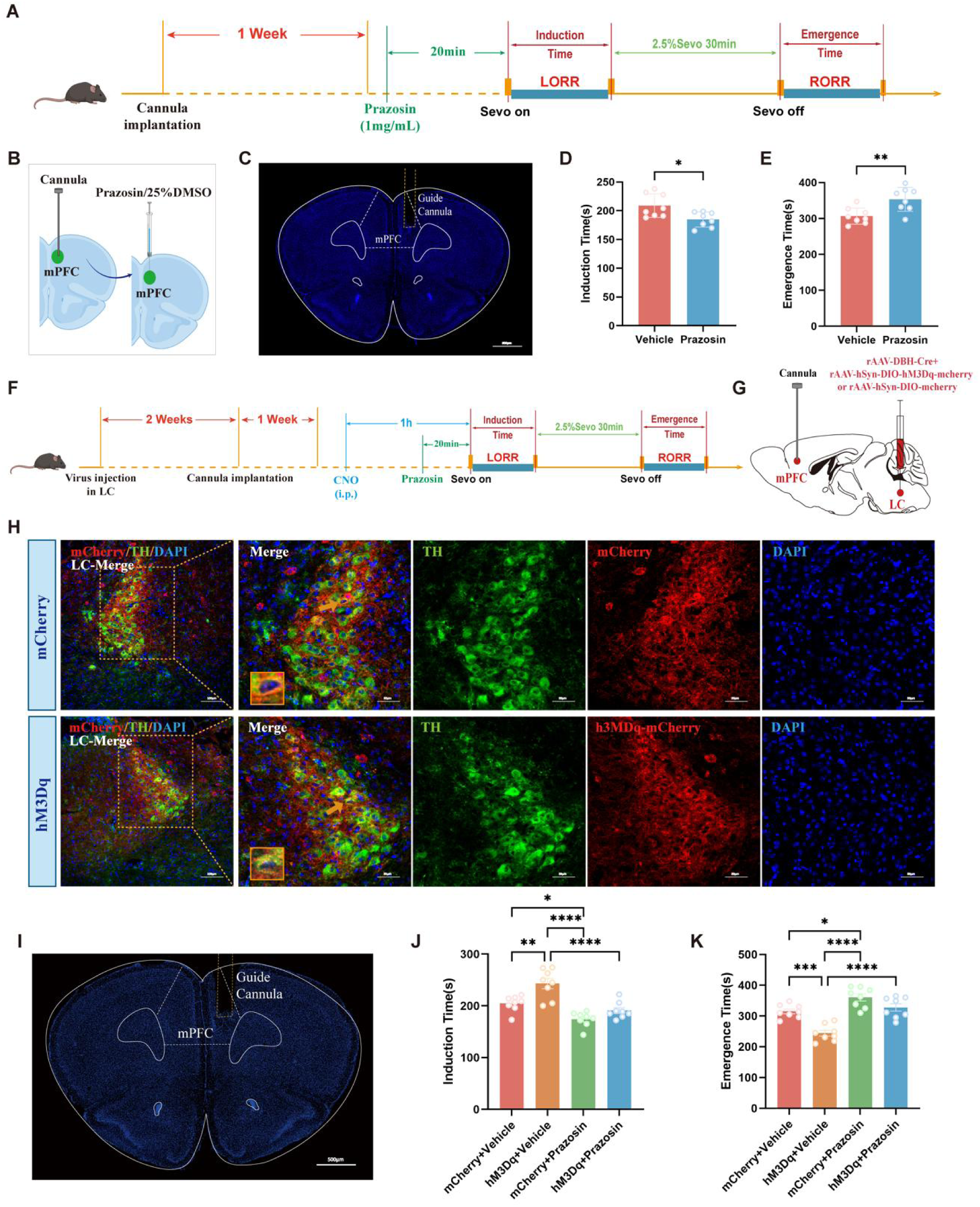
The pro-arousal effects of chemogenetic activation of LC-mPFC NEergic neural circuit can be reversed by α1-AR antagonist. **(A)** Experimental protocol for intra-mPFC microinjection of 1mg/ml α1-AR antagonist Prazosin. **(B)** Schematic of the position of the microinjection of Prazosin in the mPFC. **(C)** Representative image of the track of cannula implanted into the mPFC. **(D, E)** Induction time and emergence time for intra-mPFC microinjection in vehicle and Prazosin groups. **(F)** Experimental protocol for chemogenetic activation of LC^NE^ neurons and intra-mPFC microinjection of α1-AR antagonist Prazosin. **(G)** Schematic of the position of the intra-mPFC microinjection of Prazosin and the virus injection in the LC. **(H)** Representative images show the location of virus injection in the LC and the co-expression of hM3Dq-mCherry/mCherry, TH and DAPI in mCherry and hM3Dq groups. **(I)** Representative image of the track of cannula implanted into the mPFC. **(J, K)** Induction time and emergence time for chemogenetic activation in mCherry + Vehicle, hM3Dq + Vehicle, mCherry + Prazosin and hM3Dq + Prazosin groups. ****p<0.0001; ***p<0.001; **p<0.01; *p<0.05; i.p., Intraperitoneal injection; Sevo, Sevoflurane; LORR, Loss of Righting Reflex; RORR, Recovery of Righting Reflex

### 3.8 The GABAergic and NEergic systems interact with each other and jointly regulate the consciousness transition from sevoflurane

We implanted cannulas into the locus coeruleus of mice to monitor the changes in calcium signaling within this brain region following the injection of the GABAA-R antagonist gabazine (Fig. 8A-B). We confirmed that the cannulas implantation position was accurate in the LC (Fig. 8C) and the fluorescent images revealed a significant increase in c-Fos expression in LC^NE^ neurons after gabazine antagonize GABAA-R (P<0.0001, Fig. 8D-E).We found that microinjection of 5 μg/mL gabazine into the locus coeruleus significantly reduced the recovery time compared to saline (P<0.01, Fig. 8F). Additionally, gabazine also significantly increased the proportion of c-fos(+)TH(+)/TH(+) cells in the locus coeruleus (P<0.0001, Fig. 8G). These findings suggest that the GABAA-R in the locus coeruleus might be the initial binding site where sevoflurane exerts its effects on altering consciousness. To investigate the role of GABAA-R in the regulation of sevoflurane recovery by locus coeruleus neurons, we used shRNA to knock down the expression of GABAA-R in neurons of the bilateral locus coeruleus. Upon assessment via immunofluorescence staining analysis, we found that the expression of GABAA-R in locus coeruleus neurons was significantly reduced compared to the sham-operated group (P<0.0001, Fig. 8H-O). In our previous experiments, we initially demonstrated through pharmacological approaches that the GABAergic and NE systems jointly regulate recovery from sevoflurane. Subsequently, we combined genetic knockdown techniques with fiber photometry to conduct more in-depth investigations (Fig. 9A-F). We found that, compared to the sham-operated group, the peak calcium signal ΔF/F in the NEergic terminals of the mPFC was significantly reduced in the shRNA group (P<0.01, Fig. 9G-I).

**Figure 8.**
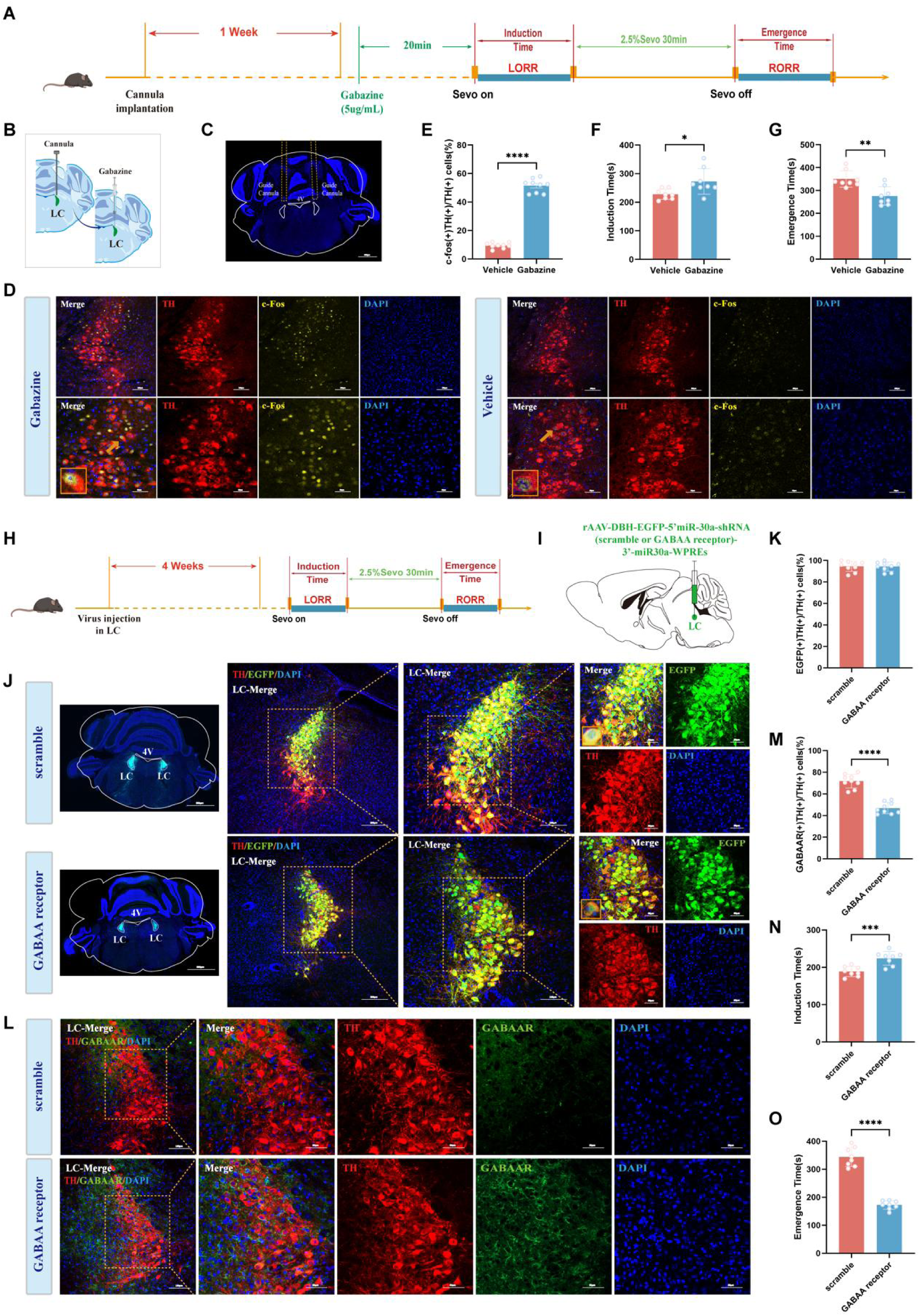
Effects of intra-LC microinjection of gabazine and knockdown of GABAA-R on the induction time of and emergence time from sevoflurane anesthesia. **(A)** Experimental protocol for intra-LC microinjection of 5mg/ml GABA-AR antagonist Gabazine. **(B)** Schematic of the position of the microinjection of Gabazine in the LC. **(C)** Representative image of the track of cannula implanted into the LC. **(D)** Representative images show the co-expression of and the staining for c-Fos, TH and DAPI in Gabazine and Vehicle groups. **(E)** The quantification of c-fos(+) TH(+)/TH(+) cells in the LC in Vehicle and Gabazine groups. **(F, G)** Induction time and emergence time for intra-LC microinjection in Vehicle and Gabazine groups. **(H)** Experimental protocol for knockdown of GABAA-R in the LC. **(I)** Schematic of the position of the virus injection in the LC. **(J)** Representative coronal brain slice and images show the location of virus injection in the LC and the co-expression of EGFP, TH and DAPI in scramble and GABAA receptor groups. **(K)** The quantification of EGFP(+)TH(+)/TH(+) cells in the LC in scramble and GABAA receptor groups. **(L)** Representative coronal brain slice and images show the co-expression of and the staining for GABAAR, TH and DAPI in scramble and GABAA receptor groups. **(M)** The quantification of GABAAR(+) TH(+)/TH(+) cells in the LC in scramble and GABAA receptor groups. **(N, O)** Induction time and emergence time for knockdown of GABAA-R in scramble and GABAA receptor groups. ****p<0.0001; ***p<0.001; **p<0.01; *p<0.05; Sevo, Sevoflurane; LORR, Loss of Righting Reflex; RORR, Recovery of Righting Reflex

**Figure 9.**
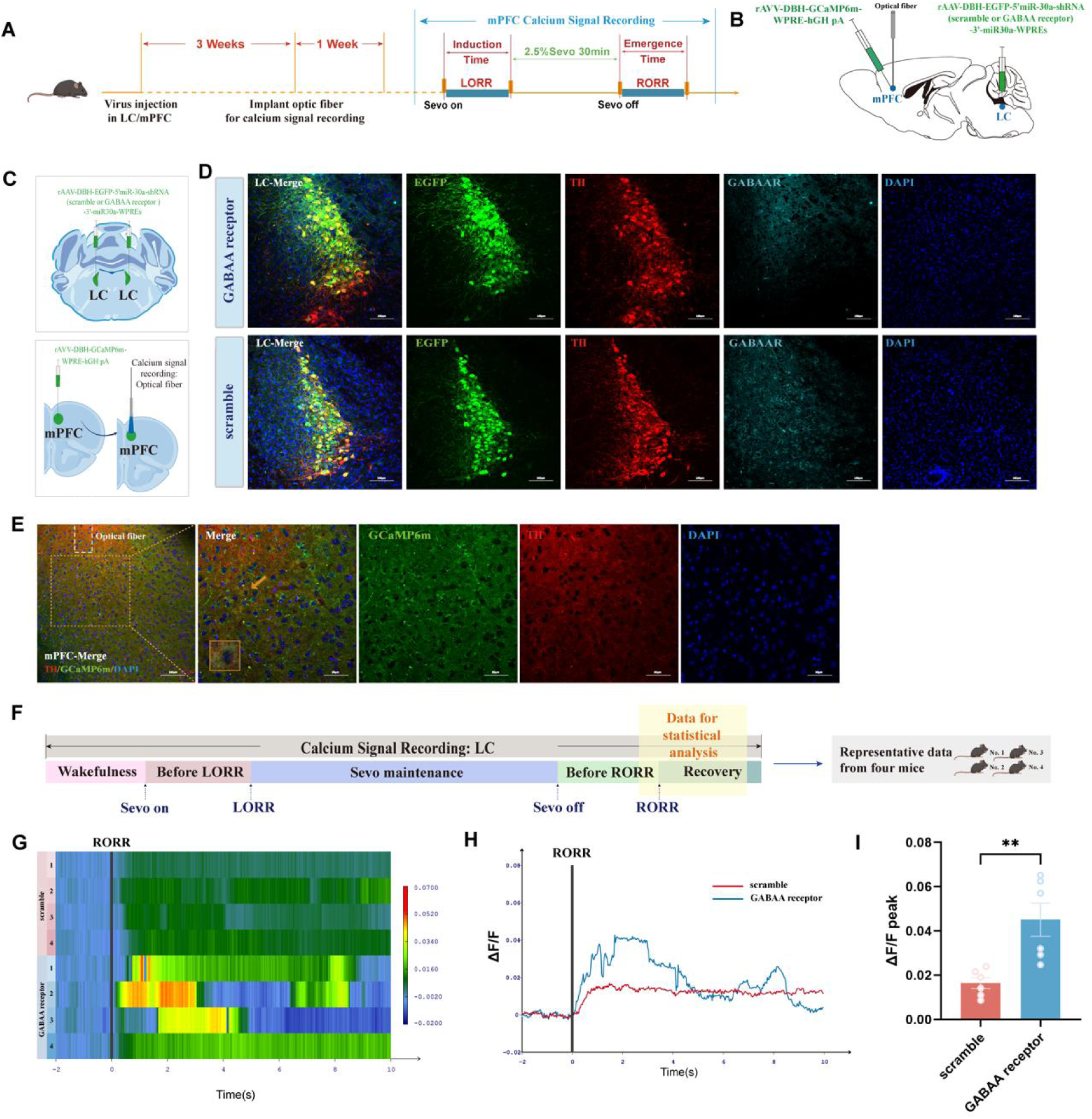
GABAA-R is an important mechanical binding site for sevoflurane anesthesia in the LC. **(A)** Experimental protocol for GABAA-R knockdown and calcium signal recording of mPFC NE neurons. **(B, C)** Schematic of the position of virus injection in the LC and mPFC and optic fibers implantation in the mPFC. **(D)** Representative images show the location of virus injected in the LC and the co-expression of and the staining for GABAAR, EGFP, TH and DAPI in scramble and GABAA receptor groups. **(E)** Representative images, showing the location of virus injected in the mPFC, and representative images of staining for DAPI, TH and GCaMP6m in the mPFC. **(F)** Schematic diagram of the different stages of the calcium signal recording in the LC and the sources of statistical data. **(G-I)** Heatmap and statistical diagram show the activity wave of calcium signals and the peak ΔF/F in the mPFC for GABAAR knockdown in scramble and GABAA receptor groups during the RORR stage. **p<0.01; Sevo, Sevoflurane; LORR, Loss of Righting Reflex; RORR, Recovery of Righting Reflex

## 4 Discussion

The specific mechanisms underlying the reversible loss of consciousness induced by general anesthesia remain a focal point of research in the field of anesthesiology. Due to the complexity of these mechanisms, despite preliminary progress in current research, many critical questions have yet to be fully elucidated. To address this gap, we selected sevoflurane, a commonly used anesthetic in clinical practice, and conducted studies in mice to identify a promising target and develop an effective strategy that offers new insights into facilitating the transition of consciousness during sevoflurane anesthesia-awakening, thereby expanding the investigation of neural network mechanisms involved in general anesthesia.

In this study, we first employed quantitative analysis of c-Fos immunofluorescence staining to verify the pivotal role of the central NE system in the transition of consciousness during sevoflurane anesthesia-awakening. We initially observed a decrease in the activity of LC^NE^ neurons during sevoflurane anesthesia. Secondly, based on calcium signaling, we found that the activity of LC^NE^ neurons was significantly inhibited during sevoflurane anesthesia but gradually recovered after anesthesia cessation. Subsequently, through pharmacological approaches, we discovered that IP injection of Atomoxetine, an NE reuptake inhibitor, not only delayed the induction of sevoflurane anesthesia and promoted post-anesthesia awakening but also significantly enhanced the activity of LC^NE^ neurons. Conversely, peripheral injection of DSP-4 reversed the pro-recovery effect of Atomoxetine. Furthermore, utilizing chemogenetic and optogenetic techniques, we revealed that activating LC^NE^ neurons prolonged the induction time of sevoflurane anesthesia and shortened the recovery time, while inhibiting LC^NE^ neurons had the opposite effect. Next, we shifted our focus to the downstream pathways of the LC. Through bidirectional tracing, we identified the mPFC as a crucial downstream target of the LC. Chemogenetic and optogenetic activation of LC^NE^ neurons influenced the activity of the mPFC. Additionally, microinjection of Prazosin, an α1-AR antagonist, into the mPFC nucleus reversed the pro-awakening effect mediated by the activation of LC^NE^ neurons, suggesting that LC^NE^ neurons may regulate the excitability of the downstream mPFC brain region via α1-AR. Furthermore, we found that injection of Gabazine, a GABAA-R antagonist, into the LC nucleus shortened the recovery time from sevoflurane anesthesia, indicating that GABAA-R in the LC could be the initial target of sevoflurane. Finally, combining gene knockdown with fiber recording, we observed that knocking down the expression of LC GABAA-R using shRNA significantly reduced the peak calcium signal in mPFC NE nerve terminals, suggesting an interaction between the GABAergic and NE systems in regulating recovery from sevoflurane anesthesia. The GABAA-R on LC^NE^ neurons and the NE α1-AR on the mPFC represent key points in this process.

Over the past few decades, the altered state of consciousness due to the action of sedative drugs on molecular targets in the CNS has been a significant focus of research. GABAA-R is a well-established target for various sedatives. As a ligand-gated Cl-channel, GABAA-R mediates most rapid inhibitory neurotransmission in the central nervous system, playing a crucial role in regulating the balance between excitation and inhibition in the brain.^28^ The inhalational general anesthetic sevoflurane, similar to most general anesthetics, primarily targets GABAA-R. Studies have shown that sevoflurane has a high affinity for GABAA-R containing the δ subunit, leading to neuronal hyperpolarization and overall network inhibition^29^. Sevoflurane may also indirectly inhibit glutamatergic neurons via GABAergic interneurons, reducing activity in arousal-related brain regions such as the thalamus and cortex^30^. Based on our findings, sevoflurane initially modulates downstream neural pathways by acting on GABAA-R in the LC. We call for future research on membrane proteins and ion channels in LC or non-LC^NE^ neurons to fully understand the molecular and neural circuit targets of general anesthetics.

Previous research on the mechanisms of general anesthesia has predominantly focused on molecular targets^31, 32^. Studies have demonstrated that sevoflurane can influence a variety of ion channels and neurotransmitter systems, including GABAA-R, to alter neuronal excitability and synaptic transmission ^33–35^. However, while this multi-target mode of action can explain its extensive neuroinhibitory effects, it fails to fully elucidate the essential mechanisms underlying anesthesia. Given the numerous similarities between general anesthesia and natural sleep, an increasing number of researchers are paying attention to the connection between the two and attempting to decipher the neurobiological mechanisms through which anesthetic drugs target sleep-wake-related neural nuclei and circuits, thereby causing reversible changes in consciousness ^36–38^.

The maintenance of brain arousal state relies on the concerted action of multiple brain regions, with the ARAS controlling the state of arousal and the transition between sleep and wakefulness cycles ^39^. It has been reported that LC^NE^ neurons constitute a crucial component of the ARAS, playing a significant role in promoting wakefulness, maintaining arousal, and vigilance ^40^. Specifically, we hypothesize that LC^NE^ neurons and their associated circuits are involved in the transition of consciousness between sevoflurane anesthesia and arousal. In this study, LC^NE^ neurons were activated via optical stimulation and chemogenetics to rapidly induce a transition from unconsciousness to arousal^41, 42^. The LC, while receiving fiber projections from arousal-promoting nuclei, also sends extensive projections through its NE neurons to various brain regions such as the cerebral cortex, hypothalamus, and basal ganglia, coordinating the arousal state of the entire brain^43^. Initially, we employed immunofluorescence, calcium signaling, and pharmacological methods to verify the critical role of the central NE system in the transition of consciousness between sevoflurane anesthesia and arousal. Subsequently, based on the fact that the LC is the primary nucleus responsible for NE release in the brain, we further demonstrated the vital role of LC^NE^ neurons in this transition by selectively activating or inhibiting them using optogenetics and chemogenetics. In summary, this study elucidates the pivotal role of LC^NE^ neurons in the transition of consciousness between sevoflurane anesthesia and arousal through the application of pharmacological, optogenetic, and chemogenetic techniques.

One of the most crucial neural pathways is the projection from the LC to the cerebral cortex. Given that the mPFC, situated downstream of the LC, plays a pivotal role in regulating sleep-wakefulness and consciousness states^16^, we focused our investigation on the LC-mPFC circuitry. Functional imaging studies have demonstrated that neural activity in the mPFC is significantly correlated with anesthesia depth, and changes in its BOLD signal accurately reflect transitions in consciousness^26^. Additionally, local inhibition of neuronal activity in the mPFC through microinjection of tetrodotoxin shortens anesthesia induction time and prolongs arousal time ^44^. These findings indicate that the mPFC also plays a significant role in reversible changes in consciousness induced by general anesthesia. Recent research suggests that NE released from LC axon terminals during NREM sleep can regulate the mPFC to control spindle waves and promote sensory arousal and micro-arousals^45^. Firstly, we demonstrated that LC may play an indispensable role in the transition of consciousness between sevoflurane anesthesia and arousal, as atomoxetine, a broad norepinephrine reuptake inhibitor in the brain, lost its effectiveness following selective lesioning of the LC. Secondly, our study identified GABAA-R in the LC as potential key targets of sevoflurane’s actions. Thirdly, we pharmacologically identified the α1-AR as a necessary target and further validated it in a more specific manner. Finally, we found that microinjection of the α1-AR antagonist prazosin into the mPFC significantly reversed the arousal-promoting effects of microinjecting GABAA-R antagonists into the LC or activating LC^NE^ neurons. Crucially, atomoxetine, a psychotropic medication used to treat attention-deficit hyperactivity disorder (ADHD), emerges as a promising candidate for translational research aimed at preventing delayed recovery in ADHD patients due to midazolam abuse. Therefore, further exploration of the specific projection relationship between LC^NE^ neurons and the mPFC, as well as the impact of the LC-mPFC neural circuitry on the transition of consciousness between sevoflurane anesthesia and arousal, is necessary in our research.

The translational significance of a study is indeed a crucial indicator in gauging its importance. Our current research offers a novel perspective on understanding the neural mechanisms of general anesthesia, with the potential to translate into clinical guidelines for the use of sevoflurane. Furthermore, the arousal-promoting effect of activating the LC-mPFC neural circuit holds potential application value for optimizing anesthesia strategies and reducing delayed recovery from anesthesia. Notably, our study found that increasing central NE levels can delay the induction of sevoflurane anesthesia and promote post-anesthetic arousal, providing a theoretical basis for the development of novel anesthesia adjuvant drugs. Additionally, it offers a target for future drug development targeting α1-AR, which may aid in the creation of more precise anesthesia modulators with fewer side effects.

In summary, our research reveals the pivotal regulatory role of LC NE neurons and the LC-mPFC neural circuit in the transition of consciousness during sevoflurane anesthesia and arousal. It provides a lens through which to elucidate the dynamic changes in neural circuits within the brain during alterations in consciousness, offering new insights into reducing anesthesia-related complications. Moreover, these findings not only broaden the exploration of neural network mechanisms underlying general anesthesia but also lay a solid theoretical foundation for optimizing clinical anesthesia management and the development of novel anesthesia adjuvant drugs.

### Authors’ contributions

H.H.Z. designed the study. W.H.S., Y.Y., Y.X.W., X.Y.D. and Z.W.Z. performed and analyzed most of the experiments. L. L, L.Y. Gu, X.X. X, Z.Y. Z, J.X. G. helped with the analysis of the experiments and participated in the modification. All authors reviewed the manuscript.

## Acknowledgments

We thank YuDong Zhou and Yi Shen for their help in experimental design.

## Data Availability

The data supporting the findings of this study are available within the article. Data will be made available upon reasonable request.

## Declarations

### Ethics Approval

All procedures were in accordance with the National Institutes of Health Guidelines for the Care and Use of Laboratory Animals and were approved by the Animal Advisory Committee of Zhejiang University.

### Consent to Participate

Not applicable.

### Consent for Publication

Not applicable.

## 7 Supplemental materials

**Supplemental Table 1.**
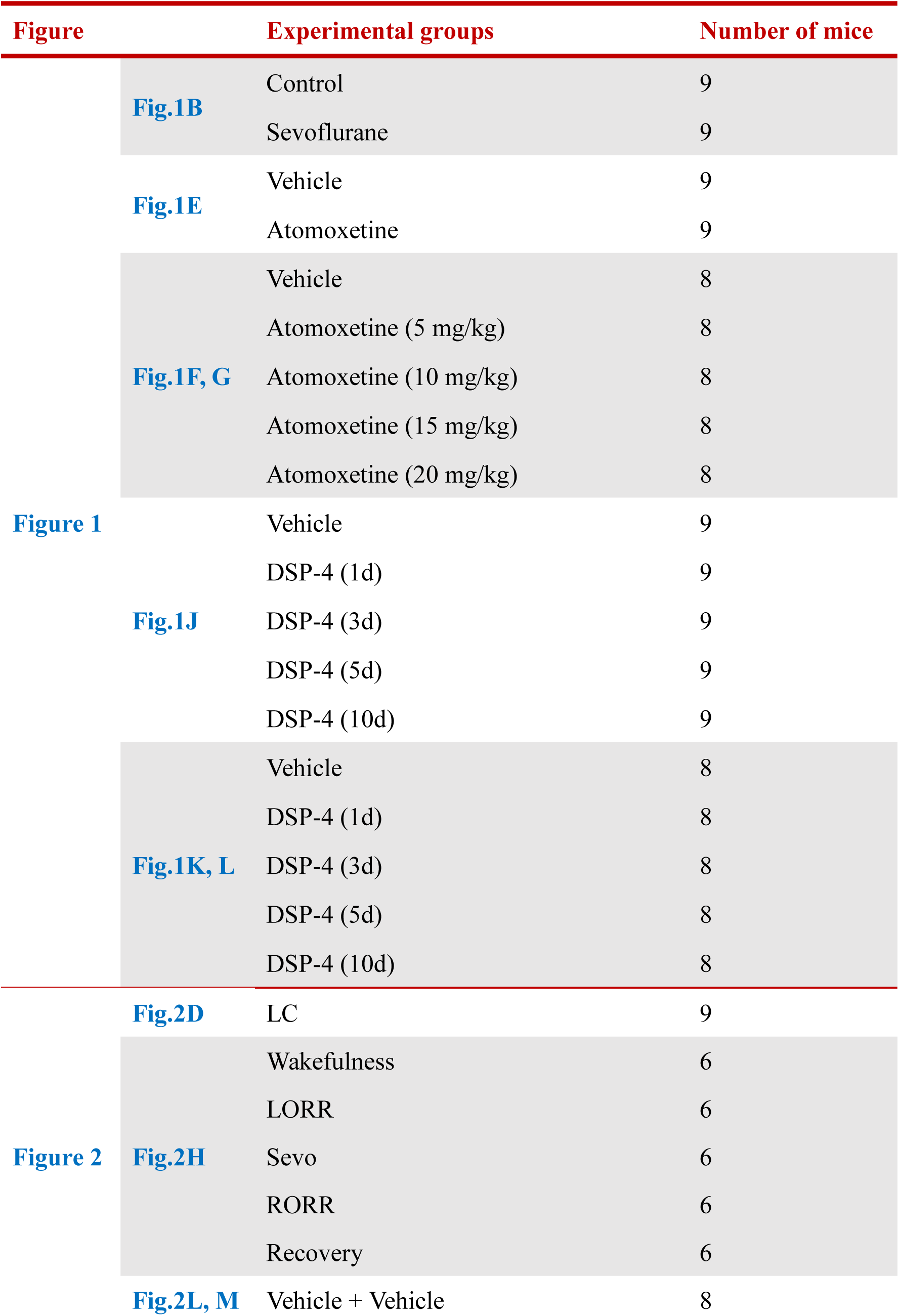
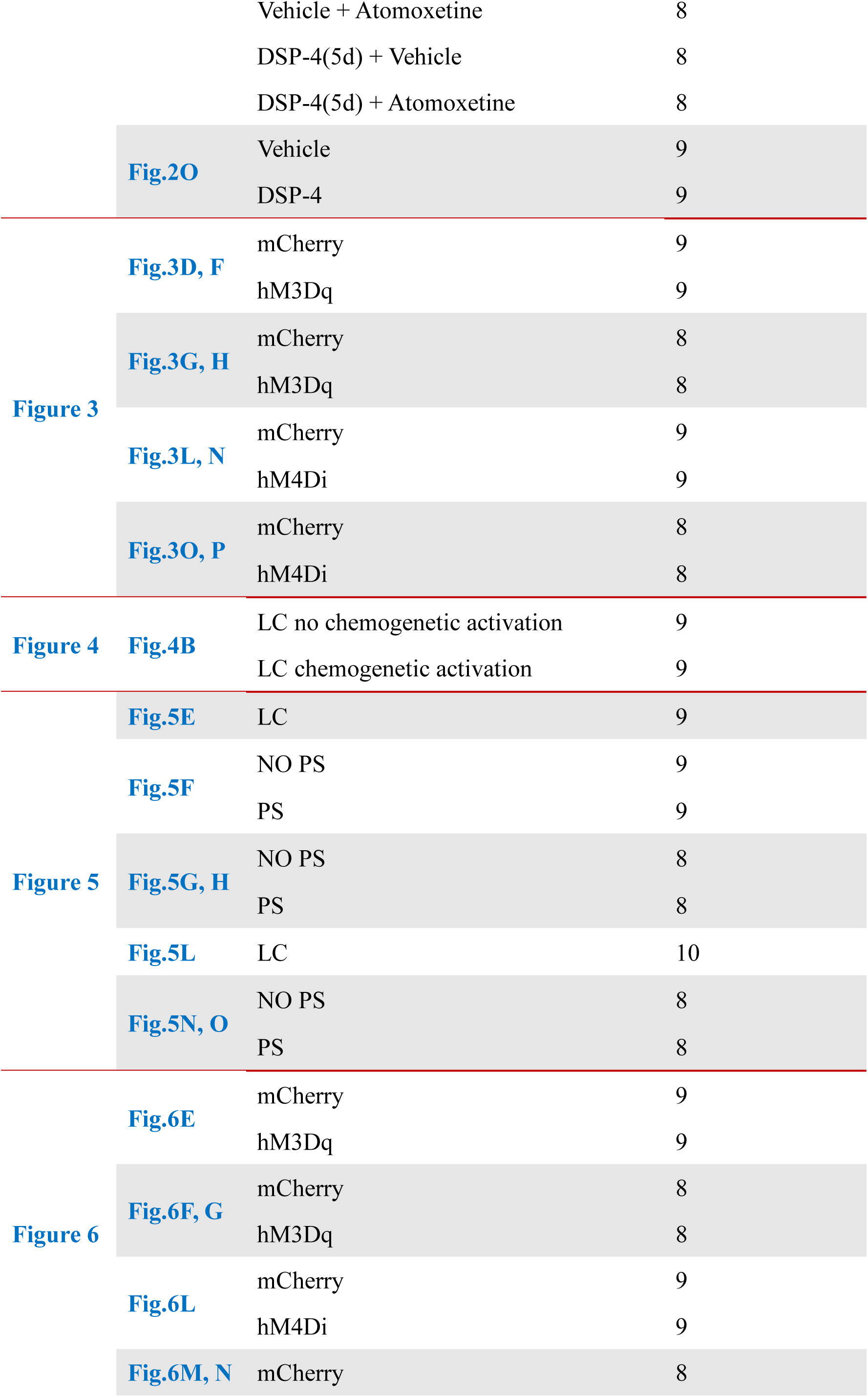
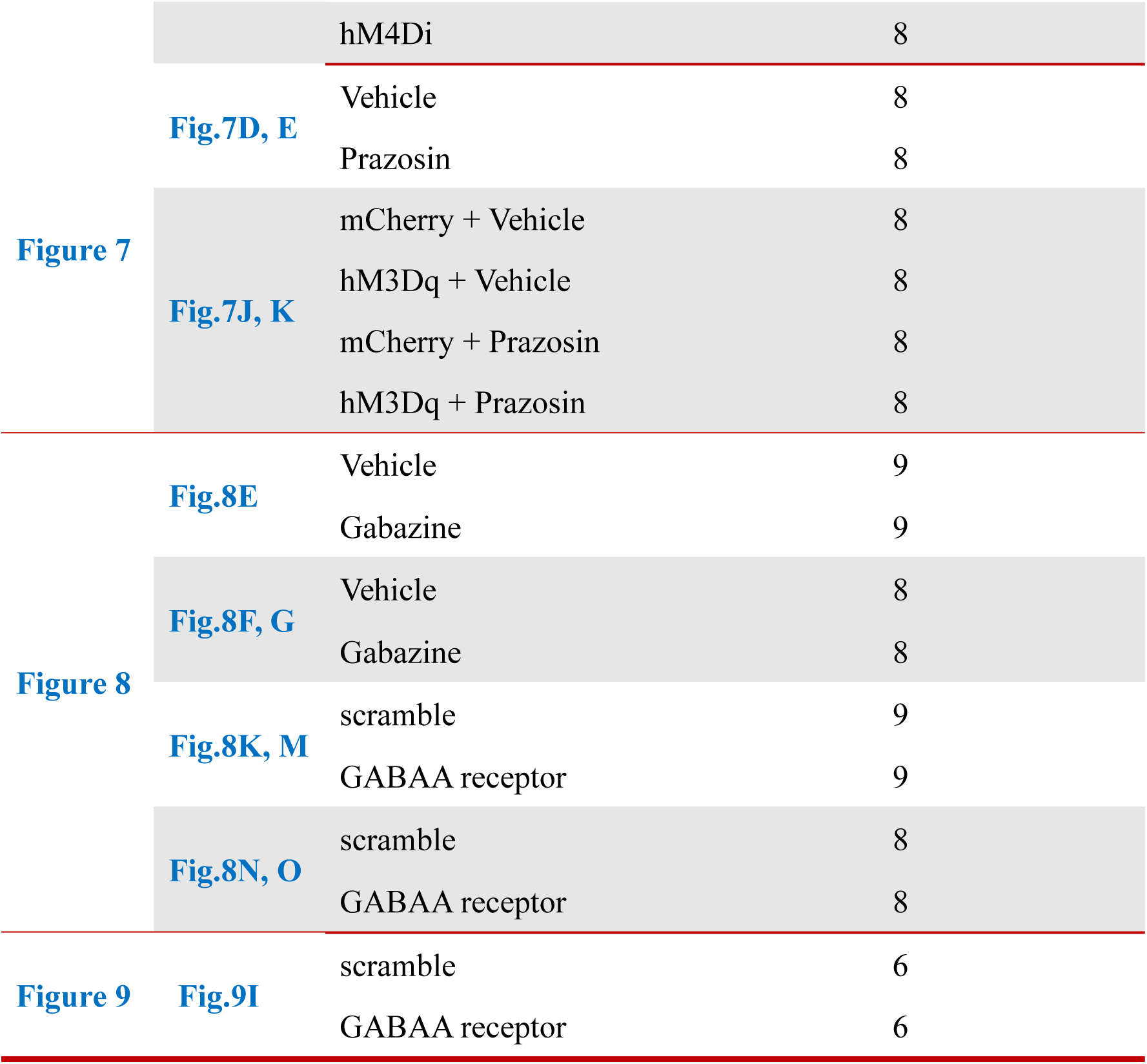
Summary of experimental groups of C57BL/6J mice.

**Table S2.**
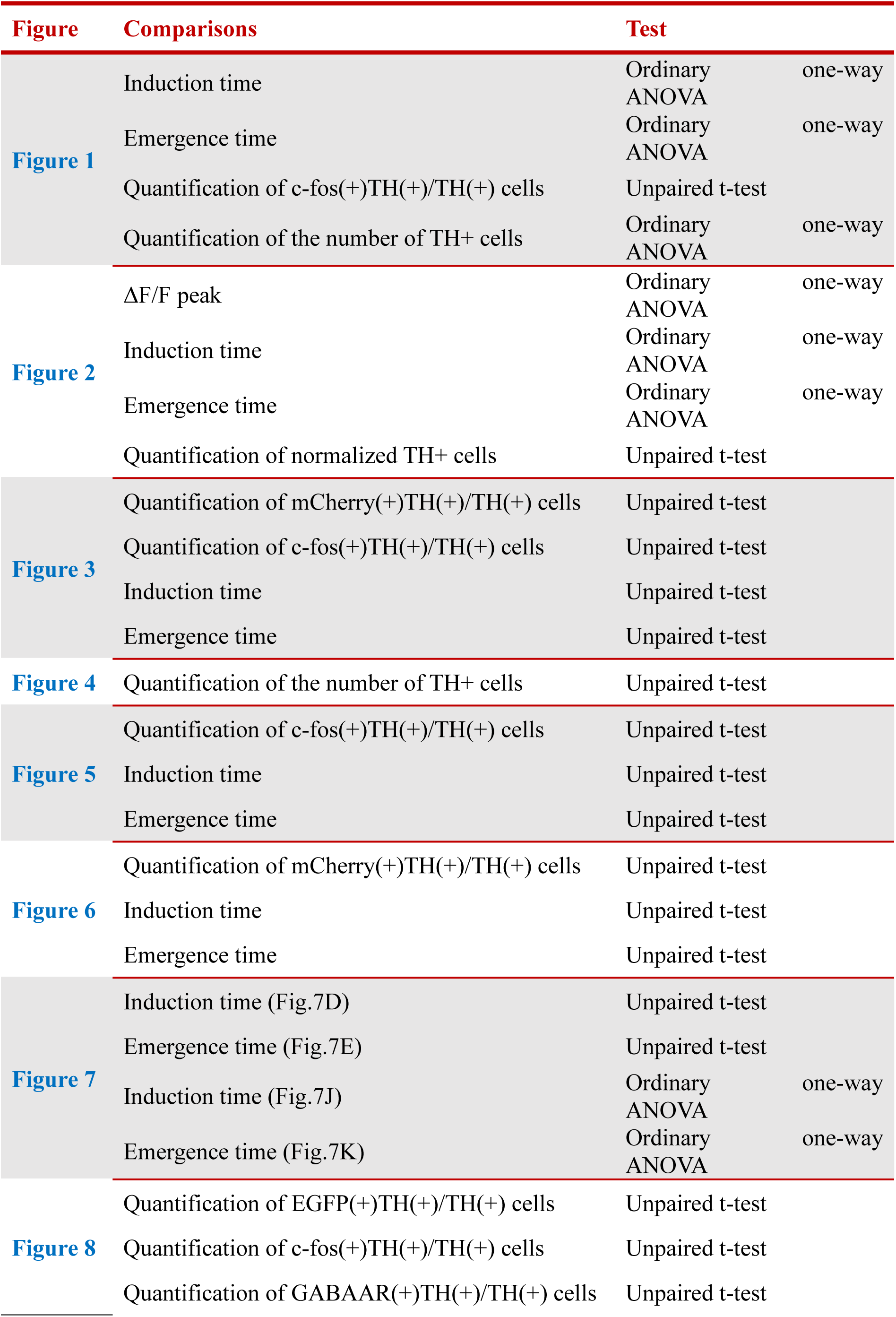
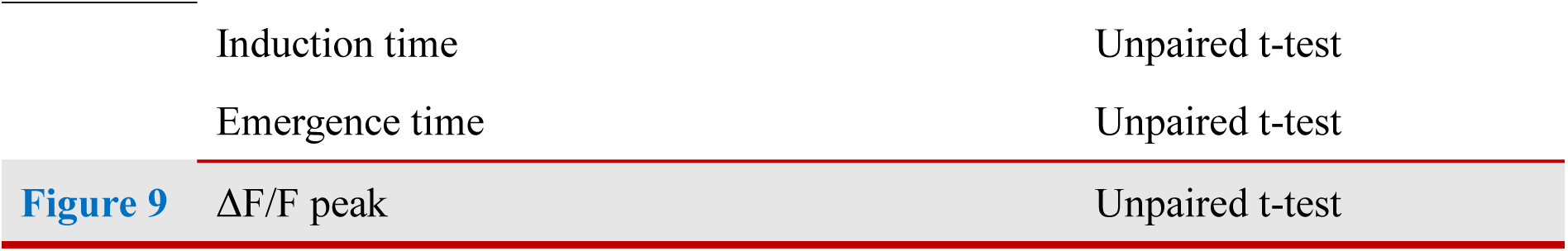
Statistical 1 analysis in each group.

**Table S3.**
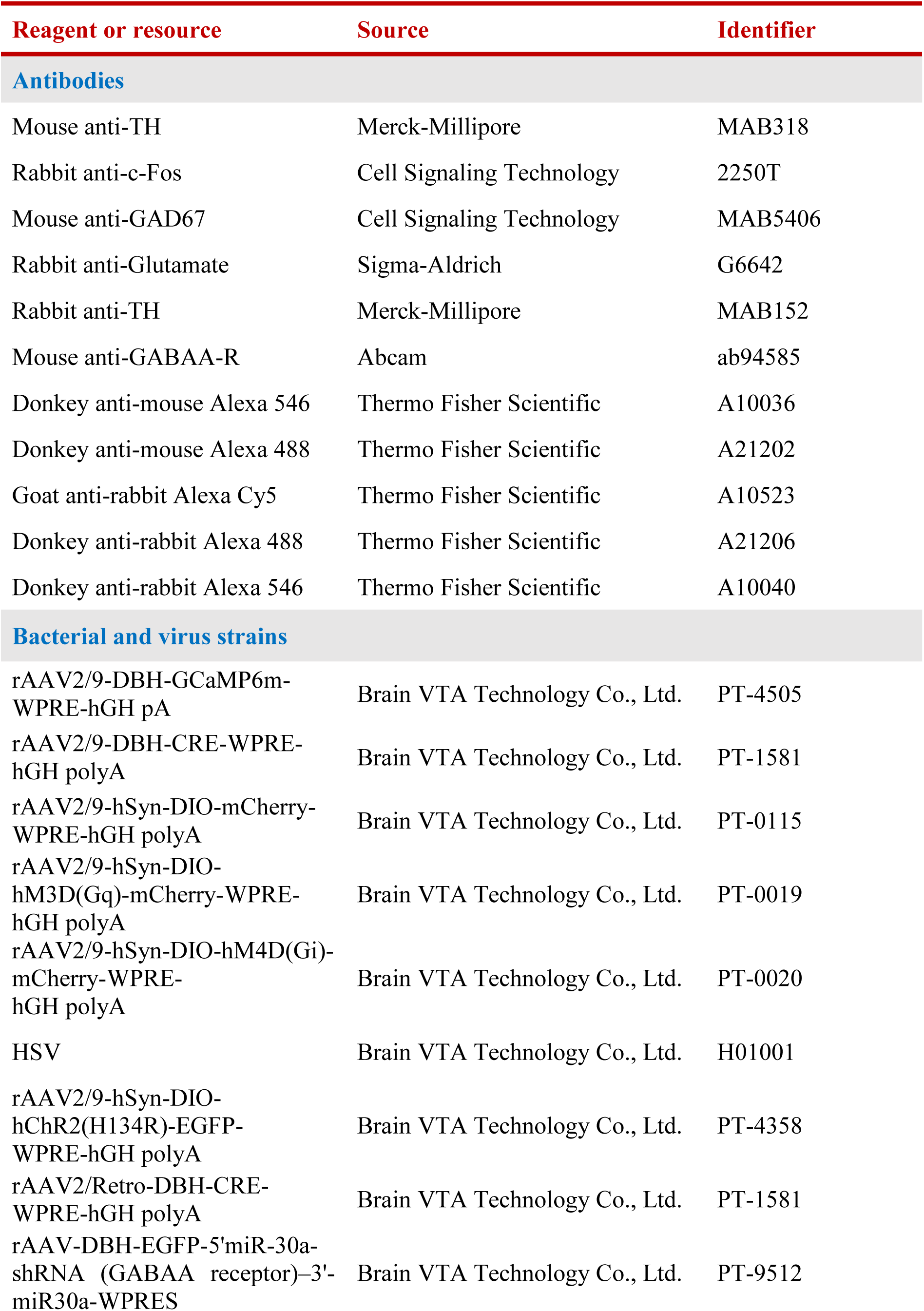
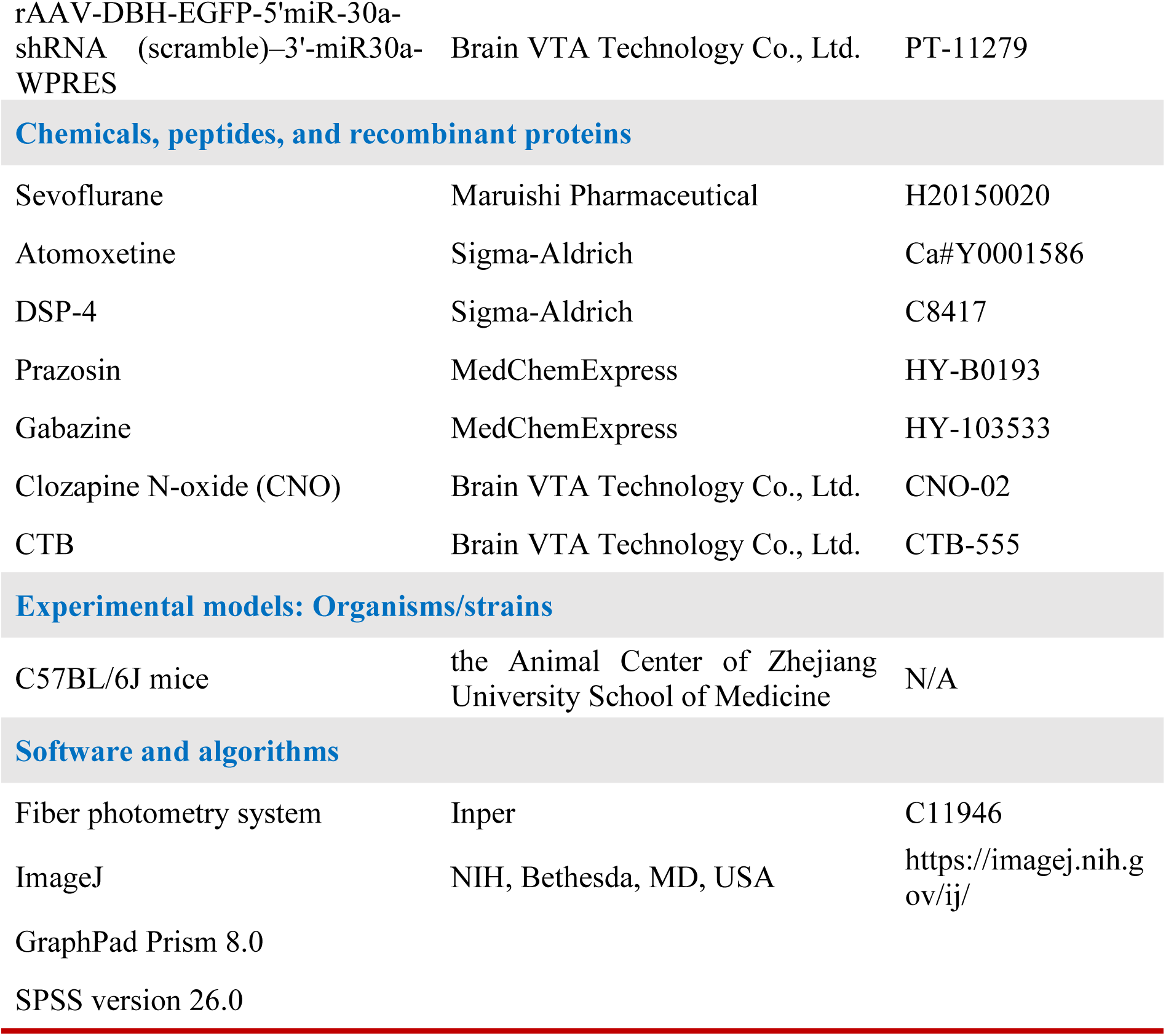
Reagent or resource.

| Reagent or resource | Source | Identifier |
| --- | --- | --- |
| **Antibodies** |  |  |
| Mouse anti-TH | Merck-Millipore | MAB318 |
| Rabbit anti-c-Fos | Cell Signaling Technology | 2250T |
| Mouse anti-GAD67 | Cell Signaling Technology | MAB5406 |
| Rabbit anti-Glutamate | Sigma-Aldrich | G6642 |
| Rabbit anti-TH | Merck-Millipore | MAB152 |
| Mouse anti-GABAA-R | Abcam | ab94585 |
| Donkey anti-mouse Alexa 546 | Thermo Fisher Scientific | A10036 |
| Donkey anti-mouse Alexa 488 | Thermo Fisher Scientific | A21202 |
| Goat anti-rabbit Alexa Cy5 | Thermo Fisher Scientific | A10523 |
| Donkey anti-rabbit Alexa 488 | Thermo Fisher Scientific | A21206 |
| Donkey anti-rabbit Alexa 546 | Thermo Fisher Scientific | A10040 |
| **Bacterial and virus strains** |  |  |
| rAAV2/9-DBH-GCaMP6m-WPRE-hGH pA | Brain VTA Technology Co., Ltd. | PT-4505 |
| rAAV2/9-DBH-CRE-WPRE-hGH polyA | Brain VTA Technology Co., Ltd. | PT-1581 |
| rAAV2/9-hSyn-DIO-mCherry-WPRE-hGH polyA | Brain VTA Technology Co., Ltd. | PT-0115 |
| rAAV2/9-hSyn-DIO-hM3D(Gq)-mCherry-WPRE-hGH polyA | Brain VTA Technology Co., Ltd. | PT-0019 |
| rAAV2/9-hSyn-DIO-hM4D(Gi)-mCherry-WPRE-hGH polyA | Brain VTA Technology Co., Ltd. | PT-0020 |
| HSV | Brain VTA Technology Co., Ltd. | H01001 |
| rAAV2/9-hSyn-DIO-hChR2(H134R)-EGFP-WPRE-hGH polyA | Brain VTA Technology Co., Ltd. | PT-4358 |
| rAAV2/Retro-DBH-CRE-WPRE-hGH polyA | Brain VTA Technology Co., Ltd. | PT-1581 |
| rAAV-DBH-EGFP-5'miR-30a-shRNA (GABAA receptor)-3'-miR30a-WPRES | Brain VTA Technology Co., Ltd. | PT-9512 |
| rAAV-DBH-EGFP-5'miR-30a-shRNA (scramble)–3'-miR30a-WPRES | Brain VTA Technology Co., Ltd. | PT-11279 |
| Chemicals, peptides, and recombinant proteins |  |  |
| Sevoflurane | Maruishi Pharmaceutical | H20150020 |
| Atomoxetine | Sigma-Aldrich | Ca#Y0001586 |
| DSP-4 | Sigma-Aldrich | C8417 |
| Prazosin | MedChemExpress | HY-B0193 |
| Gabazine | MedChemExpress | HY-103533 |
| Clozapine N-oxide (CNO) | Brain VTA Technology Co., Ltd. | CNO-02 |
| CTB | Brain VTA Technology Co., Ltd. | CTB-555 |
| Experimental models: Organisms/strains |  |  |
| C57BL/6J mice | the Animal Center of Zhejiang University School of Medicine | N/A |
| Software and algorithms |  |  |
| Fiber photometry system | Inper | C11946 |
| ImageJ | NIH, Bethesda, MD, USA | https://imagej.nih.gov/ij/ |
| GraphPad Prism 8.0 |  |  |
| SPSS version 26.0 |  |  |

## References

1. Mt A, Ag H, G T. Consciousness and anesthesia. Science (New York, NY) 2008; 322

2. D K, C N. What don’t we know? *Science (New York*, NY) 2005; 309

3. Ga M. Anesthesia and the neurobiology of consciousness. Neuron 2024; 112

4. Ww B, S J, Wm Q, Wx L, Ch M, Zl H. Understanding the Neural Mechanisms of General Anesthesia from Interaction with Sleep-Wake State: A Decade of Discovery. Pharmacological reviews 2023; 75

5. Oa M, Er Z, Kf V, et al. The Neural Circuits Underlying General Anesthesia and Sleep. Anesthesia and analgesia 2021; 132

6. Cj Y, Jl A. Inhalational anesthetics: desflurane and sevoflurane. Journal of clinical anesthesia 1995; 7

7. Ee B. Locus coeruleus. Cell and tissue research 2018; 373

8. Gr P, S F, O E, et al. Locus coeruleus: a new look at the blue spot. Nature reviews Neuroscience Nat Rev Neurosci; 2020; 21

9. G A-J, Fe B. Activity of norepinephrine-containing locus coeruleus neurons in behaving rats anticipates fluctuations in the sleep-waking cycle. The Journal of neuroscience : the official journal of the Society for Neuroscience J Neurosci; 1981; 1

10. Carter ME, Yizhar O, Chikahisa S, et al. Tuning arousal with optogenetic modulation of locus coeruleus neurons. Nat Neurosci 2010; 13: 1526–33

11. Y L, W S, A X, D H, L W, L Z. The NAergic locus coeruleus-ventrolateral preoptic area neural circuit mediates rapid arousal from sleep. Current biology : CB Curr Biol; 2021; 31

12. A L, R L, P O, et al. Dorsal raphe serotonergic neurons promote arousal from isoflurane anesthesia. CNS neuroscience & therapeutics 2021; 27

13. Tx W, B X, W X, et al. Activation of Parabrachial Nucleus Glutamatergic Neurons Accelerates Reanimation from Sevoflurane Anesthesia in Mice. Anesthesiology Anesthesiology; 2019; 130

14. T L, Ls L. Involvement of tuberomamillary histaminergic neurons in isoflurane anesthesia. Anesthesiology 2011; 115

15. Ne T, Cj VD, Jd K, et al. Optogenetic activation of dopamine neurons in the ventral tegmental area induces reanimation from general anesthesia. Proceedings of the National Academy of Sciences of the United States of America 2016; 113

16. Ga M, D P, En B. Prefrontal cortex as a key node in arousal circuitry. Trends in neurosciences 2022; 45

17. J H, De L, Kt B, S C, F W. Prefrontal cortical regulation of REM sleep. Nature neuroscience 2023; 26

18. S S, F de P, C V, et al. Disruption of posteromedial large-scale neural communication predicts recovery from coma. Neurology 2015; 85

19. A J, De N, Wh A, Jw B. Distinct Regions within Medial Prefrontal Cortex Process Pain and Cognition. The Journal of neuroscience : the official journal of the Society for Neuroscience J Neurosci; 2016; 36

20. C Z, H Z, Z N, et al. Dynamics of a disinhibitory prefrontal microcircuit in controlling social competition. Neuron Neuron; 2022; 110

21. X M, C Z, Y C, F P, Z L. Working memory and reward increase the accuracy of animal location encoding in the medial prefrontal cortex. Cerebral cortex (New York, NY : 1991) 2023; 33

22. N K, M U, Y T, et al. Activity of midbrain reticular formation and neocortex during the progression of human non-rapid eye movement sleep. The Journal of neuroscience : the official journal of the Society for Neuroscience J Neurosci; 1999; 19

23. P M. Functional neuroimaging of normal human sleep by positron emission tomography. Journal of sleep research 2000; 9

24. Aj K, Eb S, Ba M, et al. The sleep-deprived human brain. Nature reviews Neuroscience 2017; 18

25. Jb S, Z L, Gdr W, Kd A, N Z. Interhemispheric resting-state functional connectivity of the claustrum in the awake and anesthetized states. Brain structure & function Brain Struct Funct; 2017; 222

26. Y W, T Y, C Y, et al. Effects of propofol on the dopamine, metabolites and GABAA receptors in media prefrontal cortex in freely moving rats. American journal of translational research Am J Transl Res; 2016; 8

27. U L-D, M I, J L-C, et al. Molecular concentration of deoxyHb in human prefrontal cortex predicts the emergence and suppression of consciousness. NeuroImage 2014; 85 **Pt 1**

28. J B, Bg G. The Role of GABA Receptor Agonists in Anesthesia and Sedation. CNS drugs 2017; 31

29. D B, M P, Ja P, Jj L. The interaction of general anaesthetics and neurosteroids with GABA(A) and glycine receptors. Neurochemistry international 1999; 34

30. En B, Pl P, Cj VD. General anesthesia and altered states of arousal: a systems neuroscience analysis. Annual review of neuroscience 2011; 34

31. Hc H, Pm R, Mb K, et al. Towards a Comprehensive Understanding of Anesthetic Mechanisms of Action: A Decade of Discovery. Trends in pharmacological sciences 2019; 40

32. Xj S, Jj H. Neurobiological basis of emergence from anesthesia. Trends in neurosciences 2024; 47

33. Hc H, Mh A, Pa G, Jr T, Ba O, Nl H. Emerging molecular mechanisms of general anesthetic action. Trends in pharmacological sciences 2005; 26

34. J M, D G, E G, et al. The effects of the general anesthetic sevoflurane on neurotransmission: an experimental and computational study. Scientific reports 2021; 11

35. Sk O, E T, Mc S, N K, N A. Volatile anesthetic effects on isolated GABA synapses and extrasynaptic receptors. Neuropharmacology 2011; 60

36. Moody OA, Zhang ER, Vincent KF, et al. The Neural Circuits Underlying General Anesthesia and Sleep. Anesth Analg 2021; 132: 1254–64

37. Q Y, F Z, A L, H D. Neural Substrates for the Regulation of Sleep and General Anesthesia. Current neuropharmacology 2022; 20

38. Y H, Y W, L Z, M L, Y W. Neural Network Mechanisms Underlying General Anesthesia: Cortical and Subcortical Nuclei. Neuroscience bulletin 2024; 40

39. Sh L, Y D. Neuromodulation of brain states. Neuron 2012; 76

40. Ba S, S W, J T, A E-R. Neuro-orchestration of sleep and wakefulness. Nature neuroscience 2023; 26

41. Vazey EM, Aston-Jones G. Designer receptor manipulations reveal a role of the locus coeruleus noradrenergic system in isoflurane general anesthesia. Proc Natl Acad Sci U S A 2014; 111: 3859–64

42. Y A, B Y, C Z, et al. Locus Coeruleus to Paraventricular Thalamus Projections Facilitate Emergence From Isoflurane Anesthesia in Mice. Frontiers in pharmacology 2021; 12

43. La S, L L. Organization of the locus coeruleus-norepinephrine system. Current biology : CB 2015; 25

44. Er H, T G, Cw F, T L, Ga M, D P. Inactivation of Prefrontal Cortex Delays Emergence From Sevoflurane Anesthesia. Frontiers in systems neuroscience 2021; 15

45. C K, M A, N H, et al. Memory-enhancing properties of sleep depend on the oscillatory amplitude of norepinephrine. Nature neuroscience 2022; 25

